# Self-organization of embryonic stem cells into a reproducible embryo model through epigenome editing

**DOI:** 10.1101/2024.03.05.583597

**Authors:** Gerrald A. Lodewijk, Sayaka Kozuki, Clara Han, Benjamin R. Topacio, Abolfazl Zargari, Seungho Lee, Gavin Knight, Randolph Ashton, Lei S. Qi, S. Ali Shariati

**Affiliations:** Department of Biomolecular Engineering, University of California, Santa Cruz, CA; Genomics Institute, University of California, Santa Cruz, CA; Institute for The Biology of Stem Cells, University of California, Santa Cruz, CA; Department of Electrical and Computer Engineering, University of California, Santa Cruz, CA; Neurosetta LLC, Madison, WI; Wisconsin Institute for Discovery, Madison, WI; Department of Biomedical Engineering, University of Wisconsin-Madison, Madison, WI; Department of Bioengineering, Stanford University, Stanford, CA; Sarafan ChEM-H, Stanford University, Stanford, CA; Chan Zuckerberg Biohub – San Francisco, San Francisco, CA

## Abstract

Embryonic stem cells (ESCs) can self-organize *in vitro* into developmental patterns with spatial organization and molecular similarity to that of early embryonic stages. This self-organization of ESCs requires transmission of signaling cues, via addition of small molecule chemicals or recombinant proteins, to induce distinct embryonic cellular fates and subsequent assembly into structures that can mimic aspects of early embryonic development. During natural embryonic development, different embryonic cell types co-develop together, where each cell type expresses specific fate-inducing transcription factors through activation of non-coding regulatory elements and interactions with neighboring cells. However, previous studies have not fully explored the possibility of engineering endogenous regulatory elements to shape self-organization of ESCs into spatially-ordered embryo models. Here, we hypothesized that cell-intrinsic activation of a minimum number of such endogenous regulatory elements is sufficient to self-organize ESCs into early embryonic models. Our results show that CRISPR-based activation (CRISPRa) of only two endogenous regulatory elements in the genome of pluripotent stem cells is sufficient to generate embryonic patterns that show spatial and molecular resemblance to that of pre-gastrulation mouse embryonic development. Quantitative single-cell live fluorescent imaging showed that the emergence of spatially-ordered embryonic patterns happens through the intrinsic induction of cell fate that leads to an orchestrated collective cellular motion. Based on these results, we propose a straightforward approach to efficiently form 3D embryo models through intrinsic CRISPRa-based epigenome editing and independent of external signaling cues. CRISPRa-Programmed Embryo Models (CPEMs) show highly consistent composition of major embryonic cell types that are spatially-organized, with nearly 80% of the structures forming an embryonic cavity. Single cell transcriptomics confirmed the presence of main embryonic cell types in CPEMs with transcriptional similarity to pre-gastrulation mouse embryos and revealed novel signaling communication links between different embryonic cell types. Our findings offer a programmable embryo model and demonstrate that minimum intrinsic epigenome editing is sufficient to self-organize ESCs into highly consistent pre-gastrulation embryo models

## Introduction

New frontiers are emerging in the study of mammalian embryogenesis through engineering stem cell-based embryo models (SEMs) using the unrestricted potential of embryonic stem cells and their self-organizing behavior [1–23]. The first generation of SEMs can reproduce key aspects of spatial organization of pre- and post-gastrulation stages of embryonic development with potential to implant [2,24] or advance to later stages of development [15]. SEMs have the potential to transform our understanding of principles of development and self-organization of the first stages of embryogenesis and how they may go awry during catastrophic developmental abnormalities.

Current methods for generation of SEMs rely on some or all of the following: (i) Stochastic differentiation of ESCs [1,2,15,25], (ii) Biochemical optimization of media with morphogens and small molecule inhibitors or activators [2,9,13] for different embryonic cell types, and (iii) Exogenous overexpression of transcription factors [3,5,10,13,18]. During embryogenesis, however, epigenetic changes in the endogenous regulatory elements of cell fate-determining factors drives differentiation of specialized cell types that co-develop into an embryo. We hypothesized that epigenetic recapitulation of endogenous regulatory elements of key cell fate-determining transcription factors is sufficient to form major embryonic cell types and self-organize them into spatially-ordered embryo-like structures. By leveraging the programmability of CRISPR epigenome editing tools, we showed that activation of just two endogenous non-coding regulatory elements of cell fate-determining transcription factors, Cdx2 and Gata6, is sufficient to self-organize mouse embryonic stem cells into embryo-like structures with organized spatial patterns and cavity formation reminiscent of the mouse embryos at day 5-6 post-fertilization [26]. Based on these findings, we present a straightforward approach to generate highly consistent mouse embryo models in a controlled manner through intrinsic epigenetic control and without dependency on externally-added signaling factors. CRISPR activation (CRISPRa)-induced cells co-develop into embryo-like models, termed CPEMs, within three days in a simple medium with nutrients and serum, simply through chemical induction of CRISPRa. We use single-cell fluorescence time-lapse microscopy to reveal a fascinating collective cellular motion behavior that results in distinct positioning of cells in spatially-organized embryonic patterns. Single-cell transcriptome analysis of CPEMs showed clear emergence of distinct embryonic cell types expressing markers of each lineage while capturing known, as well as potential new, signaling pathway interactions between embryonic cell types. In sum, we propose a CRISPRa-based method to self-organize embryonic stem cells into mouse embryo models with highly consistent spatial organization in a facile, controlled manner without requiring external signaling cues. Our CRISPRa-based approach will pave the way for future straightforward systematic CRISPR-based screening of embryo models to enhance our understanding of epigenetic requirements for self-organization of embryonic patterns and decoding signaling networks in early embryo development.

## Results

### Generation of CRISPRa-induced primitive endoderm-like cells and trophoblast-like cells

Exogenous overexpression of coding sequences of cell fate-determining transcription factors has been widely used for controlled differentiation or reprogramming of cells. However, chromatin context, presence of co-factors, excessive expression level and epigenetic barriers can limit the activity of exogenous transcription factors. We hypothesized that direct CRISPR activation of endogenous regulatory elements can overcome epigenetic barriers associated with differentiation of embryonic stem cells [27–30]. To test our hypothesis, we implemented a CRISPRa-based strategy to generate major cell types that comprise the preimplantation embryo: epiblast stem cells (Epi), trophoblast stem cells (TS), and primitive endoderm cells (PrE), also known as hypoblast cells in human development. CRISPRa can be used to directly and efficiently activate regulatory elements of cell fate-determining transcription factors to induce robust and homogeneous lineage differentiation in stem cells. We generated a doxycycline-inducible CRISPRa embryonic stem cell line to activate endogenous regulatory elements of two major cell fate-determining transcription factors, Cdx2 (sgCdx2-mESCs) and Gata6 (sgGata6-mESCs), that are expressed in TS cells and PrE cells respectively [10,31–35].

As a control, we used mESCs expressing a non-targeting control sgRNA (sgCtrl-mESCs), as well as cells that were not treated with doxycycline (**Figure 1A**). We designed our sgRNAs to recruit CRISPRa to proximal regulatory elements of Gata6 and Cdx2 that show increased chromatin accessibility during natural development of primitive endoderm and trophoblast cells in mouse embryos [36,37] (**Supplemental Figure 1A, Supplemental Table 1)**. 24 hours after induction of CRISPRa in sgGata6-mESCs grown in a simple medium with serum and nutrients, we observed a robust and specific expression of Gata6 which was sustained for three days. Importantly, induction of Gata6 was followed by expression of two additional canonical markers of primitive endoderm, Foxa1 and Sox17, and a rapid reduction in the level of Pou5f1(Oct4) which is a marker of pluripotency (**Figure 1B-C, Supplemental Figure 1B**). In sgCtrl-mESCs or sgGata6-mESCs without doxycycline treatment, we did not observe major increases in expression of Gata6 and the primitive endoderm markers, Foxa1 and Sox17, or a decrease of the pluripotency factor Pou5f1 (**Supplemental Figure 1B**). Similar to sgGata6-mESCs, we observed robust and specific expression of Cdx2 in sgCdx2-mESCs 24 hours after induction of CRISPRa, which was not the case for sgCtrl-mESCs (**Figure 1D-E**). Notably, induction of Cdx2 was followed by expression of additional mature TE lineage markers, Tfap2c and Gata3, at day 3 and day 4 respectively, while expression of the pluripotency factor, Pou5f1, gradually declined at day 4 (**Supplemental Figure 1C**). We observed expression of Cdx2 in a percentage of sgCdx2-mESCs in the absence of doxycycline, likely because of minor leakiness of tet-responsive elements, however this was not sufficient to induce expression of mature TE lineage markers, Tfap2c and Gata3, or to decrease Oct4 expression (**Supplemental Figure 1C**). Together, these findings show that CRISPRa-mediated epigenome activation generates major cell types of the preimplantation embryo efficiently in a simple medium composed of nutrients and serum without addition of external signaling cues or exogenous expression of transcription factors.

**Figure 1.**
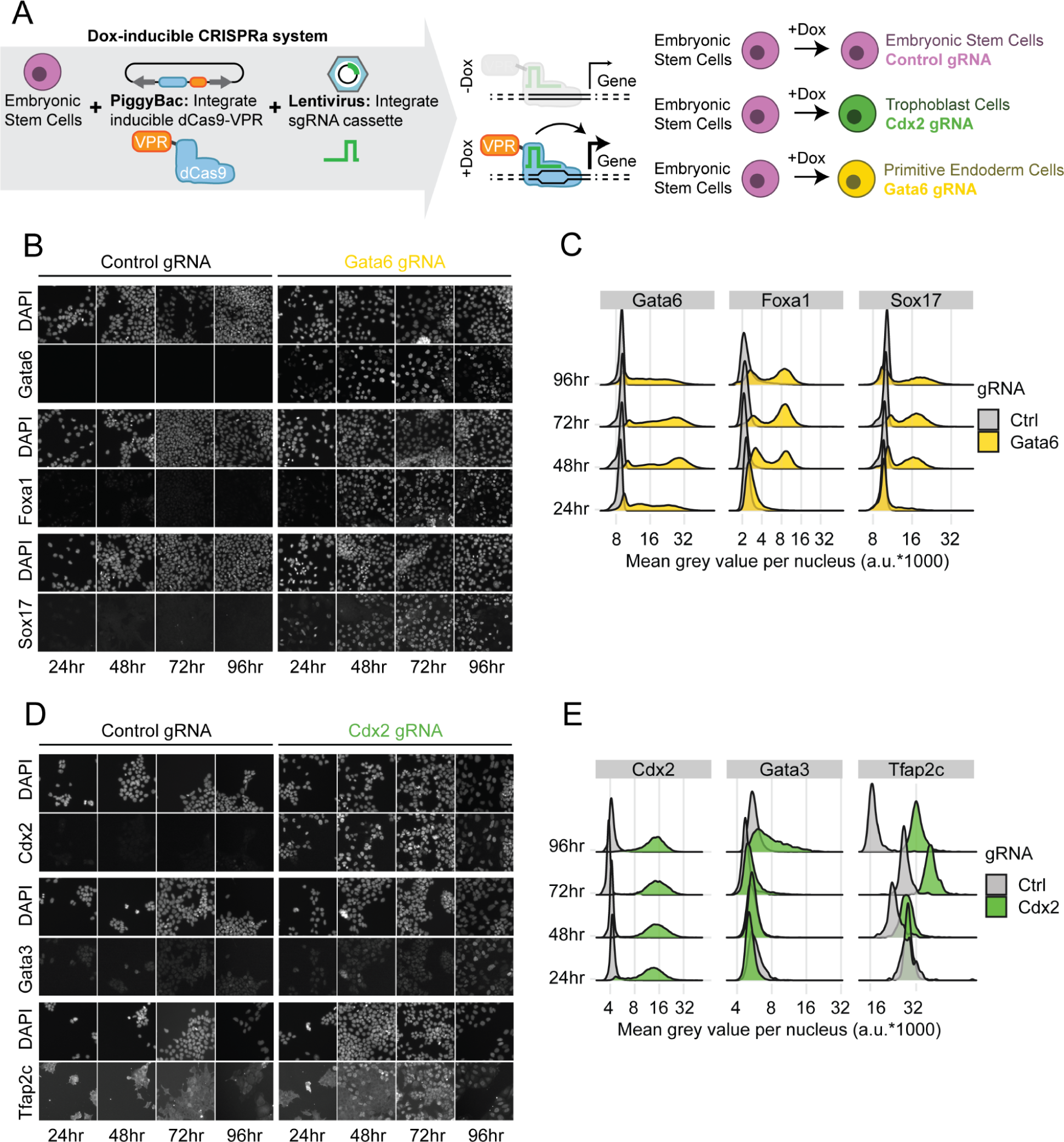
Design and validation of doxycycline-inducible CRISPRa mESCs. **A)** Overview of plasmids and integration methods. **B)** Immunofluorescence stainings of Gata6, Foxa1 and Sox17 for comparison of control and Gata6 gRNA mESCs treated with doxycycline for up to 4 days. **C)** Quantification of Gata6 induction and related PrE-cell markers over time. **D)** Immunofluorescence stainings of Cdx2, Gata3 and Tfap2c for comparison of control and Cdx2 gRNA mESCs treated with doxycycline for up to 4 days. **E)** Quantification of Cdx2 induction and related TS-cell markers over time.

### CRISPRa of 2 regulatory elements can self-organize ESCs into 2D embryonic patterns

Changes in the activity of endogenous regulatory elements are linked to differentiation of embryonic cells and emergence of spatially-organized patterns during development. We hypothesized that activation of minimum key developmental regulatory elements is sufficient to form spatially-ordered embryonic patterns. To test this hypothesis, we plated an equal mix of sgGata6-mESCs, sgCdx2-mESCs and sgCtrl-mESCs on circular micropatterns to form colonies with reproducible sizes [8,38] followed by induction of CRISPRa for 72 hours (**Figure 2A**). Next, we stained for Foxa1 to detect CRISPRa-induced sgGata6-mESCs, Cdx2 to detect CRISPRa-induced sgCdx2-mESCs and Pou5f1(Oct4) to detect sgCtrl-mESCs. CRISPRa-induced cells on circular micropatterns formed ordered structures where sgGata6-mESCs (Foxa1+) frequently formed a ring-like structure around the colony while sgCdx2-induced cells (Cdx2+) were clustered inside the colony adjacent to sgCtrl-mESCs (Pou5f1+) (**Figure 2B, Supplemental Figure 2A-B**). Interestingly, emergence of spatially-ordered 2D patterns depends on the colony size. Smaller colonies with 80 or 140 micron diameter more consistently formed a ring-like structure composed of CRISPRa-induced sgGata6-mESCs with sgCdx2-mESCs and sgCtrl-mESCs forming local clusters within the colonies as measured by distribution of Cdx2, Foxa1 and Pou5f1(Oct4) positive cells (**Figure 2B-C, Supplemental Figure 2A-D**). However, larger colonies with 225 or 500 micron diameter did not form such spatially-ordered patterns upon induction of CRISPRa (**Figure 2B-C, Supplemental Figure 2E-H**). None of these patterns emerged without doxycycline induction (**Supplemental Figure 2A-H**). This suggests that among differentiating cells, short range biochemical or biophysical interactions, or overall colony size and cell number regulates organization of cells into spatially-ordered domains. Single-cell analysis of cell number and ratio in micropatterned discs of different sizes showed that loss of patterning is correlated with an increased amount and ratio of Pou5f1+ cells, whereas we observe a decreased amount and ratio of Foxa1+ and Cdx2+ cells as the diameter of the colony increases (**Figure 2D-E, Supplemental Figure 2I**). Despite starting out with an equal ratio of cells in each region size at the seeding time, in the smaller 80 micron and 140 micron regions the Pou5f1+ population is restricted, likely by biophysical or biochemical signaling from neighboring cells, while in the larger 225 micron and 500 micron regions, cells initially are more dispersed allowing for an initial growth phase independent from neighboring cell cues. Together these results show that direct CRISPRa-based activation of 2 regulatory elements in ESCs promote differentiation and formation of spatially-organized patterns intrinsically without addition of external morphogens or chemicals. Pattern formation through intrinsic CRISPRa-induction of cell fate depends on short range biophysical and biochemical interactions, as well as overall size and cellular composition of embryonic structures.

**Figure 2.**
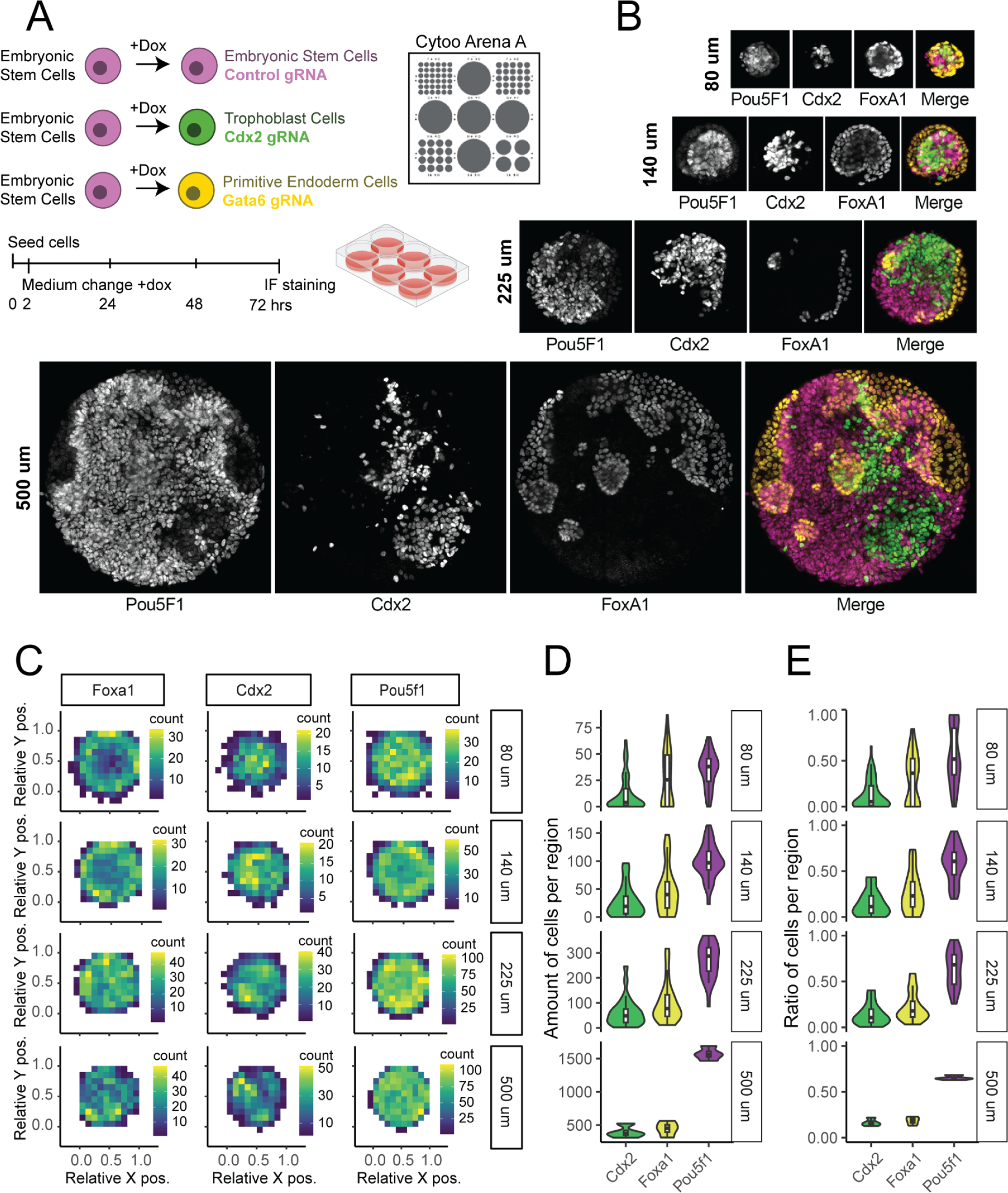
2D Micropattern organization of control-, Cdx2- and Gata6-induced mESCs. **A)** Overview of cells used and growth protocol micropattern chip. **B)** Representative immunofluorescence staining of 80um, 140um, 225um and 500um regions (top to bottom), showing markers for each mESC lines, Pou5f1, Cdx2 and Foxa1 respectively. 80um, n=62. 140um, n=35. 225um, n=25. 500um, n=4. **C)** Density plot showing average distribution of cells in normalized coordinate space for each marker and different sized regions. **D)** Distribution of total cells positive for each marker per region. **E)** Ratio of total cells positive for each marker per region.

### Collective motion of proliferating cells result in self-patterning

How do spatially-ordered embryonic patterns emerge from a few randomly positioned individual cells? To answer this question, we used single-cell live fluorescent imaging to monitor the dynamics of emergence of spatial organization in CRISPRa-programmed 2D embryonic patterns. To fluorescently visualize each CRISPRa-induced cell type, we designed new constructs in which the sgRNA for Cdx2 is co-expressed with Clover-NLS and sgRNA for Gata6 is co-expressed with mCherry-NLS (**Figure 3A**). By linking expression of each sgRNA to two different fluorescent signals, we were able to monitor both cell types in real time as they self-pattern upon addition of doxycycline or without doxycycline as a control group. After seeding the cells on RosetteArray™ well plates [39] with 100 um micropatterned regions, we imaged circular micropatterns in 1h intervals for 48 hours for both doxycycline treated cells as well as control group with no doxycycline in the media (**Figure 3A**). We used single-cell segmentation and tracking to monitor dynamics of pattern formation in 2D micropatterned discs that contained both sgCdx2-mESCs and sgGata6-mESCs, and formed ring-like structures of sgGata6-mESCs at the end of live imaging (**Figure 3B**) [40–44]. We monitored the pattern of cellular motion as cells self-organized into ordered structures. There were no discernible cellular motion patterns for the first few hours. However, we observed noticeable differences in patterns of cellular motions of CRISPRa-induced cells in later time points. CRISPRa-induced sgCdx2-mESCs cells self-assemble into confined clusters moving towards each other over time (**Figure 3B, Supplemental Movie 1A**). On the other hand, sgGata6-mESCs cells displayed a fascinating cooperative behavior by gradually moving to the edges of the micropatterns which was followed by a collective rotational movement to form a ring around other cells (**Figure 3B, Supplemental Movie 1A-B**). In contrast to CRISPRa-induced cells, non-doxycycline treated control cells moved randomly and did not form any specific pattern, as indicated by interspersed distribution of Clover and mCherry signals (**Figure 3C**). At early time points, we noticed that pattern formation begins with just a few (4-8) CRISPRa-induced sgGata6-mESCs and sgCdx2-mESCs that undergo active proliferation, albeit with different rates; CRISPRa-induced sgCdx2-mESCs displayed a slower growth rate towards the end of the 48 hour experiment, when compared to CRISPRa-induced sgGata6-mESCs or to no dox control group (**Figure 3D**). To test how induction of cell fate by CRISPRa changes migratory behavior of cells, we measured mean square displacement (MSD) over time for both sgGata6-mESCs and sgCdx2-mESCs (**Figure 3E**). We observed a noticeable increase in MSD for CRISPRa-induced sgGata6-mESCs and sgCdx2-mESC over time when compared to the no dox control group, suggesting an enhanced cellular motion upon cell fate induction and specification by CRISPRa. This shift in MSD appeared to co-occur with sgGata6-mESCs migrating to the edge at around 16-24 hours of induction, and clustering of sgCdx2-mESCs after about 32 hours of induction.

**Figure 3.**
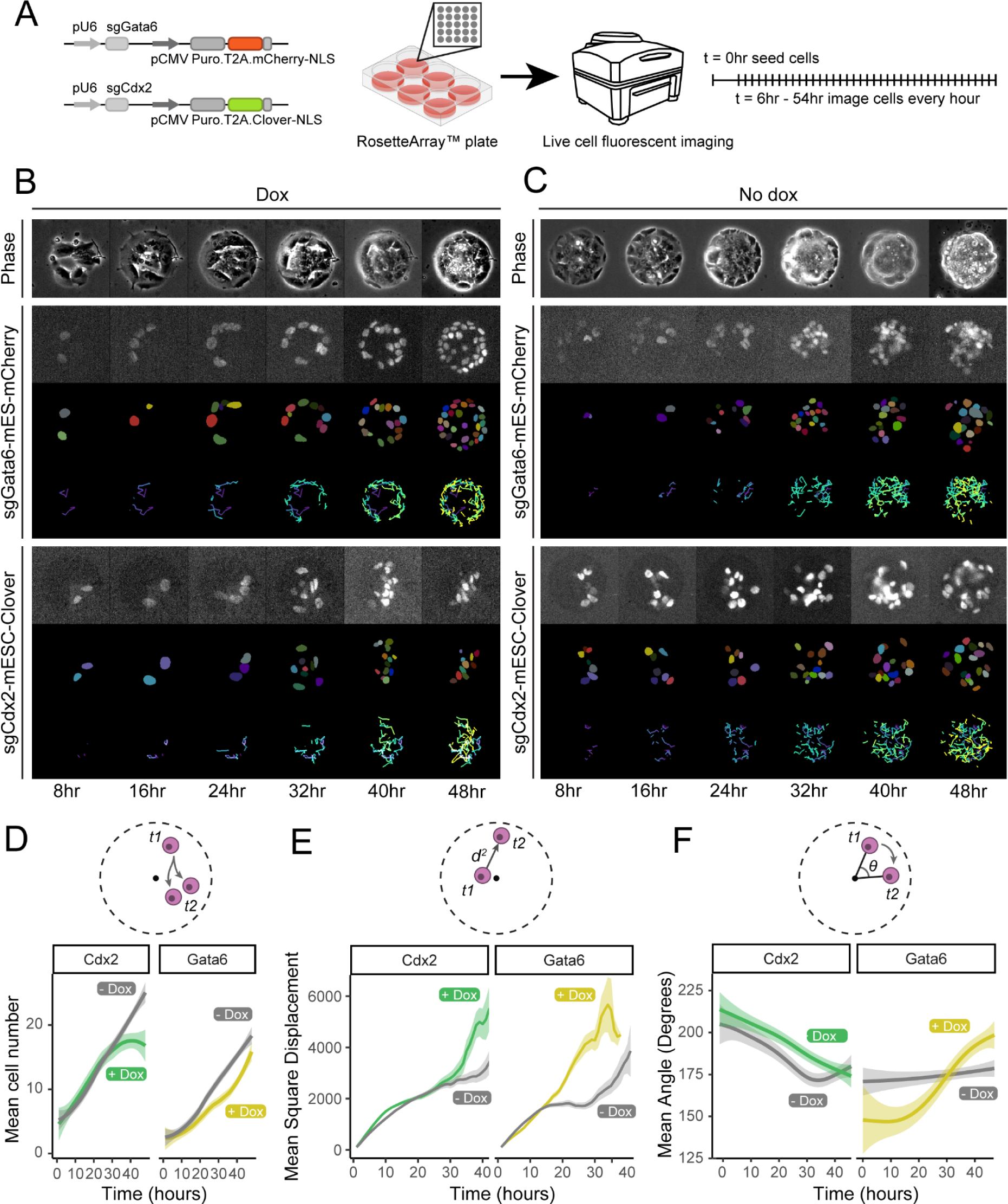
Live cell imaging and tracking of micropattern organization. **A)** Overview of modified sgRNA constructs for tracking and imaging design. **B)** Example timelapse images from a +dox (left) and **C)** -dox (right) micropattern, showing phase contrast (top), sgGata6 mESCs (middle) and sgCdx2 mESCs (bottom). For sgGata6 mESC and sgCdx2 mESC panels, top rows are fluorescent signals, middle rows are segmented nuclei, and bottom rows are cell tracking over time. **D)** Cell proliferation as measured by the number of sgCdx2-mESCs and sgGata6-mESCs over time. **E)** Average mean square displacement of sgCdx2-mESCs and sgGata6-mESCs over time. **F)** Directionality of displacements as measured by mean degree angles over time. n=11 micropattern regions per analysis. +dox sgCdx2 labels across all timepoints n=6,856, +dox sgGata6 labels across all timepoints n=3478, -dox sgCdx2 labels across all timepoints n=7,482, -dox sgGata6 labels across all timepoints n=4918.

To test if this enhanced cellular motion is paired with directionality, we quantified the angular movement of cells within individual micropattern regions. The results show that CRISPRa-induced cells display different patterns of angular movements over time when compared to non-induced sgGata6-mESCs and sgCdx2-mESCs, which was more pronounced for sgGata6-mESCs. The collective changes in movement distance and movement direction of induced sgGata6-mESCs is likely required for these cells to spatially organize themselves into ring-like structures during tissue morphogenesis (**Figure 3F, Supplemental Figure 3**) [45]. Together, our single-cell live imaging shows that intrinsic induction of fate in a few randomly positioned cells is sufficient to generate orchestrated cellular motion and growth behavior that results in formation of spatial patterns, suggesting a mostly underappreciated, yet important, role for dynamic cellular movement during mammalian embryonic pattern formation.

### Generation of 3D CRISPRa-Programmed Embryo Models (CPEMs)

Based on our findings that CRISPRa-induced embryonic cells can readily form spatially-organized 2D patterns, we reasoned that this approach can be used to more closely recapitulate natural embryonic developmental patterns by formation of 3D embryo-like structures. To this end, we plated sgGata6-mESCs, sgCdx2-mESCs and sgCtrl-mESCs in a 1:3:1 ratio in AggreWell 400 plates, which enhances consistency in size and shape of 3D cellular aggregates by confining cells into microwells with defined sizes. We performed immunostaining and confocal imaging 72 hours after CRISPRa induction to visualize the spatial organization of 3D structures (**Figure 4A**). Confocal imaging of 3D structures showed that CRISPRa-induced embryonic cells undergo self-organization to generate embryo-like 3D structures with morphological resemblance to mouse embryos at 5–6 days after fertilization (**Figure 4B, Supplemental Movie 2A**) and similar to recently reported mouse embryo models [18]. Three days after induction of CRISPRa, we observed Foxa1+ cells (sgGata6-mESCs) forming an outer layer around radially patterned Pou5f1+ cells (sgCtrl-mESCs) and clusters of self-assembled Cdx2+ cells (sgCdx2-mESCs). By segmenting 21,477 labels from single nuclei across z-positions from 30 CPEMs and plotting the average density at each z-position, we confirmed self-organization of three CRISPRa-induced cell lines into embryo-like structures with asymmetric distribution of the three cell types along the z-axis (**Figure 4C**) akin to axis formation in the egg cylinder stage of mouse embryonic development. By quantifying nuclear staining (DAPI) across the center z-position, approximating the middle of the embryo model (**Figure 4D**), we observe cavity formation in 80% of 3D structures (**Figure 4E-F**). CPEMs show consistent size of 100 micron in the z-axis and 150 micron in x-axis and y-axis (longest axis) (**Supplemental Figure 4A**). Structures show an approximately equal amount, 40%-50% each, of Foxa1+ cells and Pou5f1+ cells, and 10%-20% Cdx2+ cells (**Figure 4G-H**). Analyzing the z-axis, we observe that Pou5f1+ cells and Cdx2+ have a preference for positioning to opposing sides of the structure (**Figure 4I-J**). Despite starting out with 3 times more sgCdx2-mESCs, these cells are the least prevalent in the 3D structures, suggesting that sgCdx2-mESCs are subject to selective mechanisms that slow their growth, decrease their numbers or are delayed in their development (**Figure 4G-H**). Interestingly, quantification of cell number and ratio in embryo models with random organization (∼20% of structures) showed that they had similar cell number and ratio to those with patterned organization and cavities (**Supplemental Figure 4B**), suggesting the presence of unknown stochastic processes for spatial pattern formation through self-organization.

**Figure 4.**
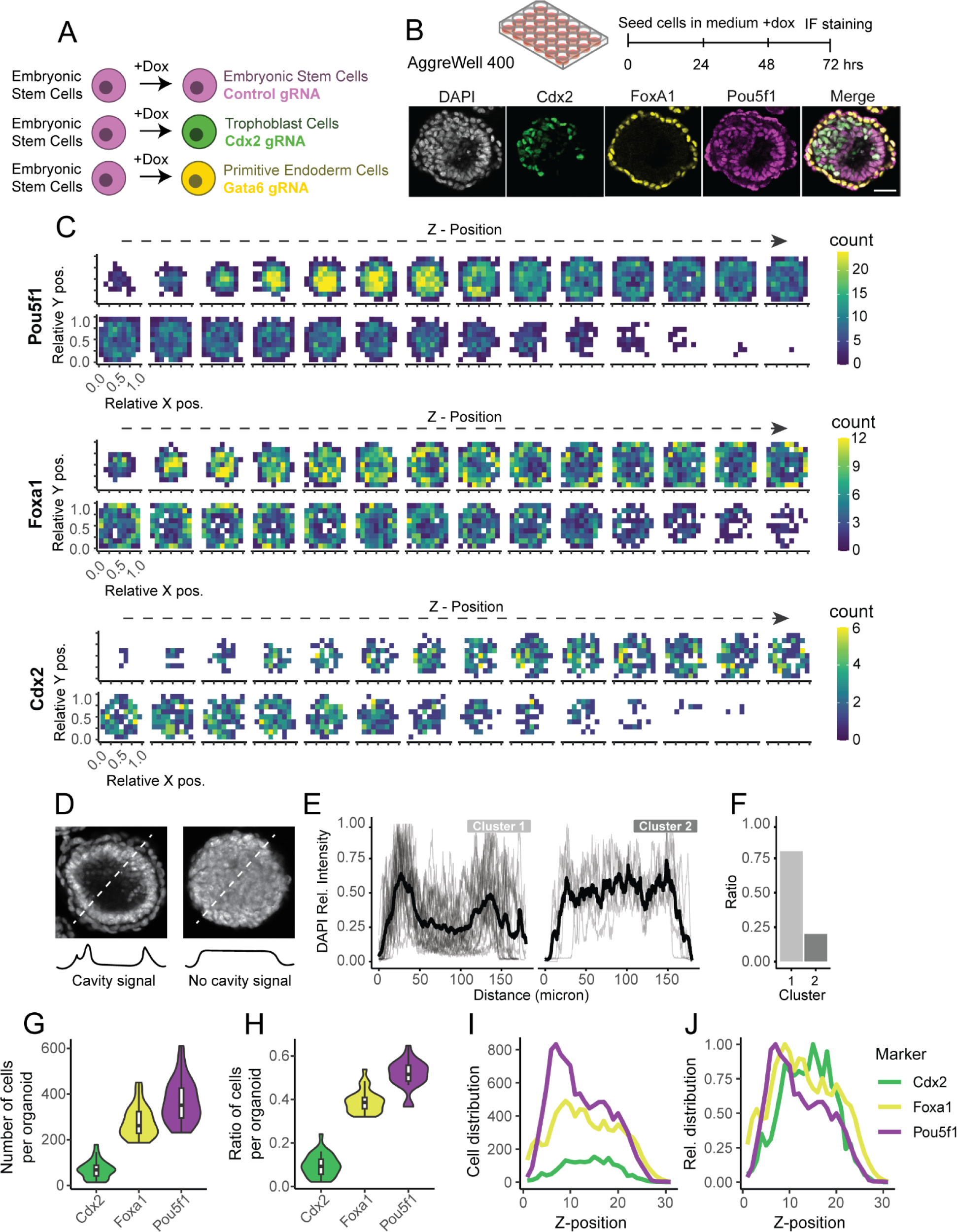
Generation and single-cell analysis of CRISPRa-programmed embryo models (CPEMs). **A)** Overview of cells used for differentiation **B)** Differentiation protocol and representative confocal image of a CPEM with each cell type present, stained by Pou5f1, Cdx2 and Foxa1. **C)** Quantification of cell density for each marker across CPEMs using z-stack confocal imaging (n=30, n=21,477 nuclei labels), left to right shows increasing depth with 3.89um z-stack step-size, 28 density maps across the z-axis are displayed. **D)** Analysis of cavity formation by analyzing DAPI stain in the center stack position of CPEMs, showing an example with (left) and without (right) cavity. **E)** Quantification and clustering of cavity formation in 2 clusters, and **F)** ratio of the clusters (n=30). **G)** Number and **H)** ratio of Cdx2, Foxa1 and Pou5f1 labeled cells identified in CPEMs (n=30). **I)** Mean number of Cdx2, Foxa1 and Pou5f1 labeled cells across imaged z-positions (n=30). **J)** Relative distribution of Cdx2, Foxa1 and Pou5f1 labeled cells across imaged z-positions (n=30), scaled to maximum within each group.

The 1:3:1 (sgCtrl-:sgCdx2-:sgGata6-mESCs) ratio of starting cells was shown to be optimal for previously published embryo models [10,46,47]. To investigate how cellular composition can affect overall spatial patterns and morphological features of CPEMs, we generated embryo models with different ratios of pre-gastrulation cell types while using models composed of only one cell type as control conditions. When using sgCtrl-mESCs alone (**Supplemental Figure 4A-B and 5A-J, Supplemental Movie 2B**) or sgCdx2-mESCs alone (**Supplemental Figure 4A-B and 6A-J, Supplemental Movie 2C**) in the 3D differentiation protocol, we did not observe any embryonic patterns. However, when using sgGata6-mESCs alone (**Supplemental Figure 4A-B and 7A-J, Supplemental Movie 2D**), we found 3D cellular structures with clear cavities, similar to cavities form in 1:3:1 embryo models, albeit with ectopic additional layering of Foxa1+ PrE-like cells on the outside of CPEMs. Moreover, when changing the ratio of cells to 1:1:3, where the sgGata6-mESCs are overrepresented, we found cavity formation is enhanced by formation of larger CPEMs with larger cavities (**Supplemental Figure 4A-B and 8A-J**). The opposite was true when changing the ratio to 3:1:1 (sgCtrl-:sgCdx2-:sgGata6-mESCs), where the sgCtrl-mESCs are overrepresented, we found additional Pou5f1+ Epi-like cells ectopically growing, filling up the cavities (**Supplemental Figure 4A-B and 9A-J**). Together, these findings show that PrE-like cells appear as essential regulators in self-patterning and cavity formation in our model.

Our quantitative single-cell maps of 3D models shows that CRISPRa-induced lines can readily and efficiently self-organize into CPEMs with spatial similarity to mouse embryos at day 5-6 post-fertilization and similar to previously reported mouse embryo models [18,46,47]. CPEMs display high consistency in cellular composition and morphological features. In addition, our findings show Gata6-expressing cells on their own can induce cavity formation, showing that these cells are likely integral to cavity formation and determining overall morphological features via interactions with other cell types in CPEMs.

### Single-cell transcriptomics uncovers major cell type and heterocellular signaling of pre-gastrulation embryonic cells

To examine the differentiation state of single cells in CRISPRa-induced cells, we performed single-cell RNA-sequencing (scRNA-Seq) on CPEMs grown from mixed cell population and, as controls, CPEMs formed from each of the CRISPRa-induced lines individually: sgCtrl-mESCs only, sgGata6-mESCs only and sgCdx2-mESCs only. Unsupervised dimensionality reduction of single-cell transcriptomics of the mixed CPEMs resolved 6 clusters that were clearly separated by expression of CRISPRa-induced target genes (**Figure 5A**). We identified 3 distinct populations as marked by Nanog, Cdx2 and Gata6 expression, corresponding to the mix of 3 cell types to generate CPEMs (**Figure 5A**). We observed two clusters of Nanog-expressing cells, annotated as Epi-1 and Epi-2, with transcriptional resemblance to embryonic epiblast cells, as shown by expression of markers such as Nanog, Pou5f1, Fgf4 and Foxd3, which likely reflect different stages of the cell cycle (**Figure 5B, Supplemental Figure 10A-C**). Gata6-expressing cells separated into 2 clusters, annotated as PrE-1 and PrE-2, which were marked by Sox17, Pdgfra, Bmp2 and Sox7 that are canonical makers of primitive endoderm (**Figure 5B, Supplemental Figure 10D**). The 2 clusters show some difference in these marker levels related to separation into parietal (Sox7, Sox17) and visceral endoderm (Bmp2, Pdgfra), suggesting the differentiation sgGata6-mESCs are subject to heterogeneity in the culture that creates the 2 subpopulations of PrE cells, possibly by their spatial positioning and differential interactions with Epi and TS cell types, or cell cycle state differences [35,48,49]. Similarly, 2 distinct populations of Cdx2-expressing cells, annotated as TS-1 and TS-2, expressed markers of trophoblast progenitors such as Tfap2c, Esrrb, Wnt5a and Lefty1 (**Figure 5B, Supplemental Figure 10E**). Cells labeled with Tfap2c and Esrrb are related to canonical differentiation of trophoblast cells, whereas the cluster with higher Wnt5a, Lefty1 and Hoxd9 expression also expressed a series of other markers likely reflecting transitory states during differentiation or a distinct subpopulation of cells [24,35,50–53]. A subset of CRISPRa-induced sgGata6-mESCs and sgCdx2-mESCs maintained the expression of pluripotency factors such as Nanog and Pou5f1(Oct4), likely due to delayed differentiation or perhaps failure to activate CRISPRa (**Figure 5B**). We also found transcriptional resemblance of the Epi, PrE and TS cell clusters to E5.25 mouse embryo scRNA-seq data [54] (**Supplemental Figure 10F-G**). The final 2 clusters, consisting of a small fraction of the total population, showed unclear association with a particular cell fate, potentially from technical artifacts in data generation, and were excluded in some of the downstream analysis. Altogether, single-cell transcriptomics data confirms successful CRISPRa induction of differentiation program of major embryonic cell types of pre-gastrulation mouse embryo, and the subsequent start of differentiation into downstream lineages.

**Figure 5.**
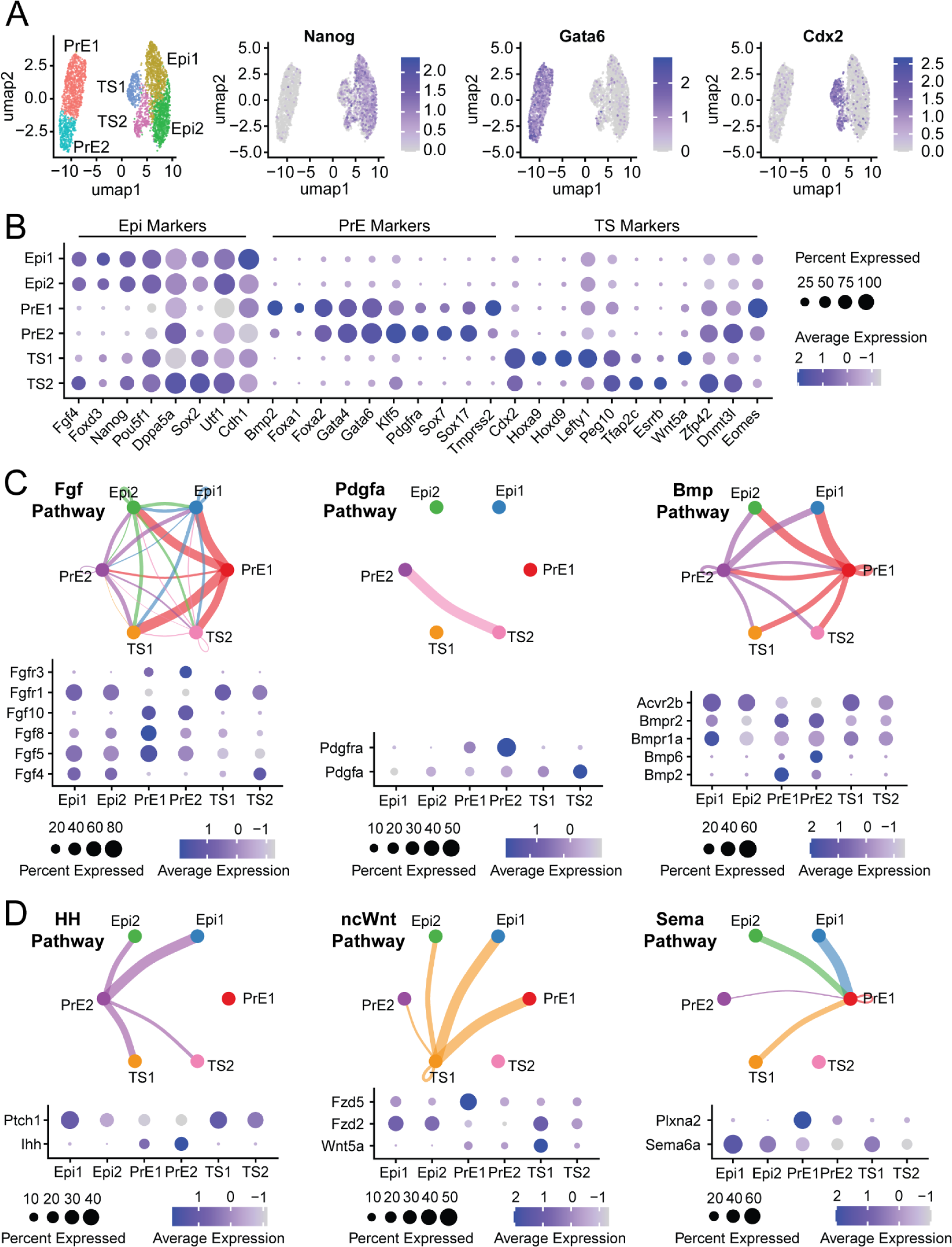
Single cell RNA-seq analysis of 3D CPEMs and ligand-receptor interactions. **A)** UMAP and clustering analysis of single cell RNA-seq data of mixed cell culture CPEMs, showing feature maps for Nanog, Gata6 and Cdx2. **B)** Dotplot showing a set of gene marker levels for Epi, TS and PrE cells. **C)** CellChat analysis showing ligand-receptor interactions between cell clusters of pathways known in early embryogenesis. **D)** CellChat analysis showing ligand-receptor interactions between cell clusters of potential new signaling communications in early embryogenesis.

To follow up on results of immunostaining data from single cell type CPEMs, we explored the transcriptional profiles of CPEMs grown from single cell types only as well. Consistently, we observed a lack of differentiation in CPEMs grown from sgCtrl-mESCs only (**Supplemental Figure 11A-B**). For sgGata6-mESCs we see most cells are Gata6 positive and a small fraction remain positive for Epiblast markers. Like the mixed CPEMs scRNA-seq data, PrE1+PrE2 show similarities to visceral endoderm and PrE3 shows similarity to parietal endoderm. (**Supplemental Figure 11C-D**). For sgCdx2-mESCs we see most cells are Cdx2 positive and a slightly larger fraction remain positive for Epiblast markers compared to the sgGata6-mESCs only, and a similar 2 TS clusters were identified compared to the mixed CPEMs data (**Supplemental Figure 11E-F**). For sgCtrl-mESC only CPEMs, the 3 Epi clusters appear to be linked to cell cycle phase again (**Supplemental Figure 12A**). Similarly, for sgGata6-mESC CPEMs, the clusters PrE1 and PrE2 are similar in transcriptional profile of marker genes, but appear different in cell cycle phase (**Supplemental Figure 12B**). No clear cell cycle phase differences were observed for the sgCdx2-mESC only CPEMs (**Supplemental Figure 12C**). Though we see global similar data for individual cell types, whether they were grown in a mixed culture or individually, small differences in cluster sizes and marker levels suggest cell-cell interactions may influence how CPEMs consisting of mixed or single cell types are shaped.

We designed our embryo model such that the three cell types start from an undifferentiated pluripotent state and co-develop into an embryo-like structure by induction of CRISPRa when seeding the cells. We also excluded the use of external stimuli delivered through recombinant proteins or small molecules in the culture medium that could otherwise bias the CRISPRa induced differentiation. We hypothesized that co-development of CRISPRa-induced cells can facilitate continuous signaling crosstalk between three cell types as they self-organize into spatially distinct embryonic patterns [20,21,55,56].

To analyze potential signaling pathway ligand-receptor interactions between clusters of different cell types, we used CellChat [57,58] to analyze paired expression of ligand-receptors between different clusters in CPEMs. We found predicted activity of pathways known to regulate early embryogenesis in specific ways, such as the Fgf pathway, Pdgf pathway and Bmp pathway [59–64] (**Figure 5C, Supplementary Figure 12D-E**). Whereas Epiblast cells are known for their secretion of Fgf4 to control differentiation of PrE and TS cells, we also find that PrE cells secrete other Fgfs including Fgf5, Fgf8 and Fgf10, which pair with a distinct Fgf receptor Fgfr3, providing a potential alternative Fgf-Mapk signaling axis originating from PrE cells and influencing Epi and TS cell development. In addition, we observed PrE cells expressing transcripts of secreted Bmp factors, Bmp2 and Bmp6, which are paired with differential expression of Bmp and Activin receptors in receiving Epi and TS cells, suggesting presence of combinatorial signaling potential in the Bmp pathway [48]. One example of a potential functional pairing is Pdgf signaling from TS cells towards PrE cells, which may promote differentiation of PrE into parietal endoderm (ParE) over visceral endoderm (VE) [35,49]. Supporting this predicted interaction, we observe a higher ratio of ParE in CPEMs grown from mixed cells (30% ParE-like) compared to sgGata6-induced cells only, which do not receive TS-cell signals (15% ParE-like) (**Supplementary Figure 12F**). We also detected paired expression of ligand-receptors in pathways that are less commonly studied in the context of early embryogenesis, such as Hedgehog, non-canonical Wnt signaling, Plexin-Semaphorin, Pecam1, Cadherin, and Ephrin pathways [65–71] (**Figure 5D, Supplementary Figure 12D-E**). Each of the pathways shown (**Figure 5D**), consisting of HH (Ihh - Ptch1), ncWnt (Wnt5a - Fzd5/Fzd2) and Semaphorin (Sema6a-Plxna2) signaling are originating from a specific clusters of PrE and TS cells, indicating they may be involved in specialized signaling centers that help pattern embryonic development. Thus, we captured multiple signaling pathways that link extraembryonic and embryonic cells, which may play an important role in differentiation and spatial organization of cells in CPEMs.

## Discussion

Like most stochastic processes, stochastic differentiation of ESCs does not allow for precise control over the starting cellular composition of different cell types that form embryo models, resulting in various degrees of variability in cellular composition and spatial arrangement. The dependency on external differentiation cues, often incompatible between different cell types, impairs co-development of different embryonic cell types into embryo models to fully capture physiological heterocellular signaling of natural embryonic development [72]. Finally, overexpression of transcription factors by adding exogenous copies of coding sequences of their genes to ESCs can allow for induction of more controlled differentiation, but may fail to overcome epigenetic barriers required for full differentiation as hypothesized by Oldak et al [9].

In addition, the induced transcription factors do not undergo natural post-transcriptional processing such as RNA splicing. As new frontiers are emerging in studying mammalian embryogenesis using stem-cell based models of embryos of different species, there is an urgent need to complement and improve existing approaches to generate embryo models in a facile and controlled manner with enhanced reproducibility in spatial patterns and cellular compositions [73].

An interesting finding of our study is that intrinsic activation of just two endogenous regulatory elements of cell fate-determining transcription factors in pluripotent cells is sufficient to self-organize cells into embryonic patterns. Based on this simple principle, we present a straightforward CRISPR-based epigenome activation method in mouse embryonic stem cells to induce different cell fates and self-organize them into embryo models. By simply mixing two cell types expressing two fate-inducing guide RNAs, Gata6, Cdx2, and a non-targeting control in a simple growth medium without addition of any external cues, we observed clear spatial organization where primitive endoderm cells (sgGata6-mESCs) are surrounding inner layers of trophoblast (sgCdx2-mESCs) and epiblast cells (sgCtrl-mESCs) with cavity formation in 3D culture models. Interestingly, we find that sgGata6-induced primitive endoderm-like cells play a major role in patterning of CPEMs, as indicated by the complete loss of patterning when these cells are not present and that sgGata6 cells on their own can generate cavitated structures, consistent with previous important role of Gata6 in cell fate decisions during early mouse embryonic development [33,64].

Our time-lapse fluorescent microscopy of 2D patterns revealed fascinating collective behavior of cells that result in emergence of self-organizing ordered structures. Active proliferation of collectively moving PrE-like results in formation of a ring-like pattern around the entire structure, whereas TS-like cells gathered together to form a local cluster inside the structure that was clearly separated from Epi-like cells. In essence, our results show that intrinsic cell fate decisions can encode molecular programs that result in formation of developmental patterns through collective motion of cells coupled with differential growth. Importantly, we showed that the size of structures influences self-organization, where PrE-like cells only reproducibly organize in smaller growth areas with 80um or 140um diameters, but not 225um or 500um. This suggests that in the right local environment, orchestrated cellular motion, growth and heterocellular signaling are sufficient to induce self-organizing embryo-like structures, without the need to additionally supply morphogens into the culture medium.

CPEMs display molecular similarities with recently published iETX and sEmbryo models, where either ectopic expression of Gata6 or Gata4, or Cdx2 are used to generate PrE or TS cells. In addition, CPEMs show similarity to iETX embryo models that have grown for 3-4 days, in both organization, cell number and cell ratio [18,46]. Specifically, day 4 iETX models were previously shown to have about 500 Epi, 150 TS and 350 PrE per structure, where we find our day 3 CPEMs contain about 350 Epi, 80 TS and 280 PrE cells each. We also find the single-cell RNAseq data from our CPEMs s can be mapped to cell types found in the E5.25 mouse embryo.

We foresee several applications for CPEMs for future experiments to investigate how functional genomic elements shape the overall morphology, cellular composition and signaling cross-talk during embryonic development. Incorporation of a programmable protein, CRISPRa, into embryo models will facilitate large-scale screening to test the role of major cell fate determining factors, enhancers and signaling factors in emergence of spatial patterns in embryo models. In addition, the inducibility of our CRISPRa device will allow for temporal control over activation of regulatory elements during embryo models. We also recognize the shortcomings of our approach that can be addressed in future application of CRISPR-based epigenome editors for generation of embryo models. The Cdx2-induced TS cells in our model appear to contribute slightly less in total number, and also may be slightly delayed in relative developmental timing, compared to Epi and PrE cells, as they maintain residual level of pluripotency factors. We anticipate that more developmentally advanced TS cells can be generated through activation of multiple regulatory elements of TS-enriched transcription factors. In line with this observation, we anticipate that targeting multiple elements can improve differentiation fidelity and timing in embryo models using multiplexable CRISPR orthologs like Cas12a [10,32,50,74,75], compared to the current single regulatory element targeting strategy. Additionally, we can use these strategies to have more control over subpopulations of cells that form, e.g. by targeting factors that for instance differentially specify parietal and visceral endoderm [49], as well as screening in a massively parallel manner for factors that induce missing cell types, such as trophoblast giant cells [76]. More precise temporal control of expression can be achieved by combining different Cas9 versions, the use of unique protein degrons [77], and the combinatorial use of specific CRISPR-activation and CRISPR-inhibition [78,79] systems to achieve highly controllable gene circuits to program cell fate decision in vitro. We anticipate implementing new generations of multiplexable, degradable CRISPR-based epigenome editors to further advance the field of stem cell-based embryo models through generation of controllable and reproducible embryo models and to overcome some of the spatiotemporal limits in the CPEM currently used in our models.

## Supporting information

Live imaging of collective cellular motion during embryo model formation

3D models of CRISPRa-progammed embryo models

## Supplemental Figures

**Supplemental Figure 1.**
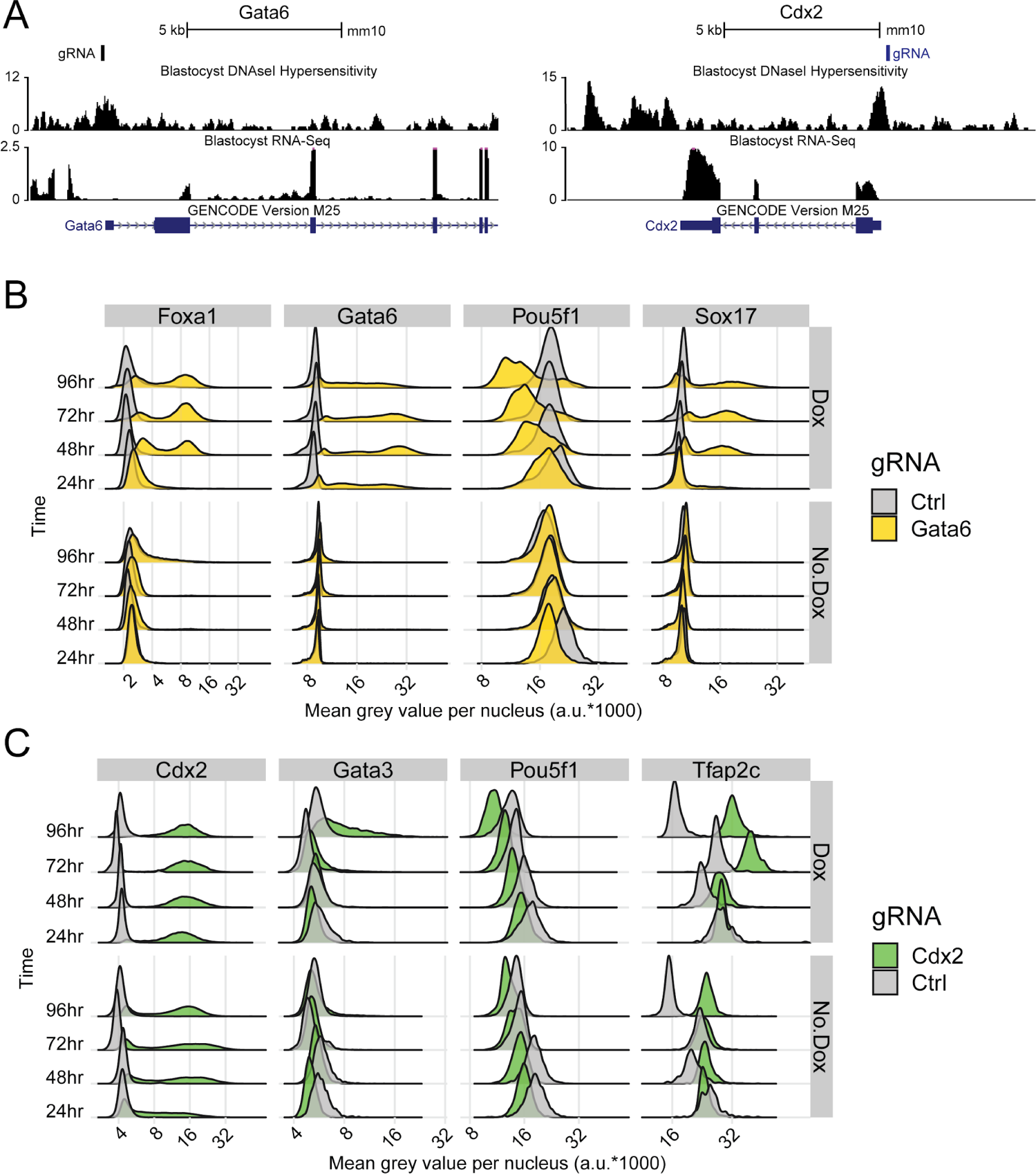
Additional validation data on CRISPRa using sgGata6. **A)** Location of Gata6 and Cdx2 gRNAs used. **B)** Quantification of CRISPRa + and - dox treatment over time, sgGata6-mESC and sgCtrl-mESCs. **C)** Quantification of CRISPRa + and - dox treatment over time, sgCdx2-mESC and sgCtrl-mESCs.

**Supplemental Figure 2.**
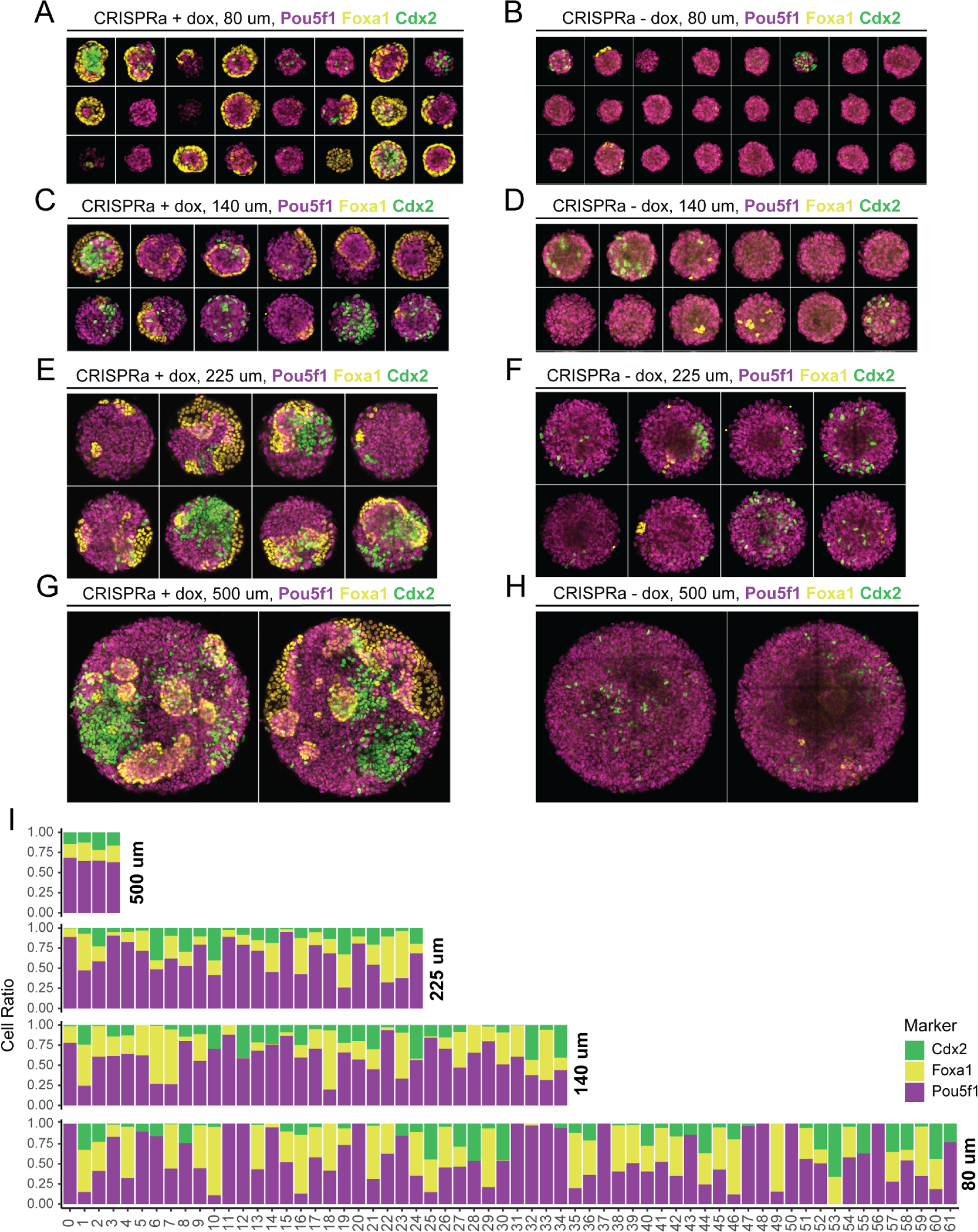
Immunostainings for doxycycline-induced CRISPRa cells grown on Cytoo micropattern chips. **A)** Example stainings for Pou5f1, Cdx2 and Foxa1 stainings of 80um +dox conditions. **B)** Example stainings for Pou5f1, Cdx2 and Foxa1 stainings of 80um -dox conditions. **C)** Example stainings for Pou5f1, Cdx2 and Foxa1 stainings of 140um +dox conditions. **D)** Example stainings for Pou5f1, Cdx2 and Foxa1 stainings of 140um -dox conditions. **E)** Example stainings for Pou5f1, Cdx2 and Foxa1 stainings of 225um +dox conditions. **F)** Example stainings for Pou5f1, Cdx2 and Foxa1 stainings of 225um -dox conditions. **G)** Example stainings for Pou5f1, Cdx2 and Foxa1 stainings of 500um +dox conditions. **H)** Example stainings for Pou5f1, Cdx2 and Foxa1 stainings of 500um -dox conditions. **I)** Ratio of cells labeled Cdx2+, Foxa1+ and Pou5f1+ for all individual micropattern regions analyzed for all sizes.

**Supplemental Figure 3.**
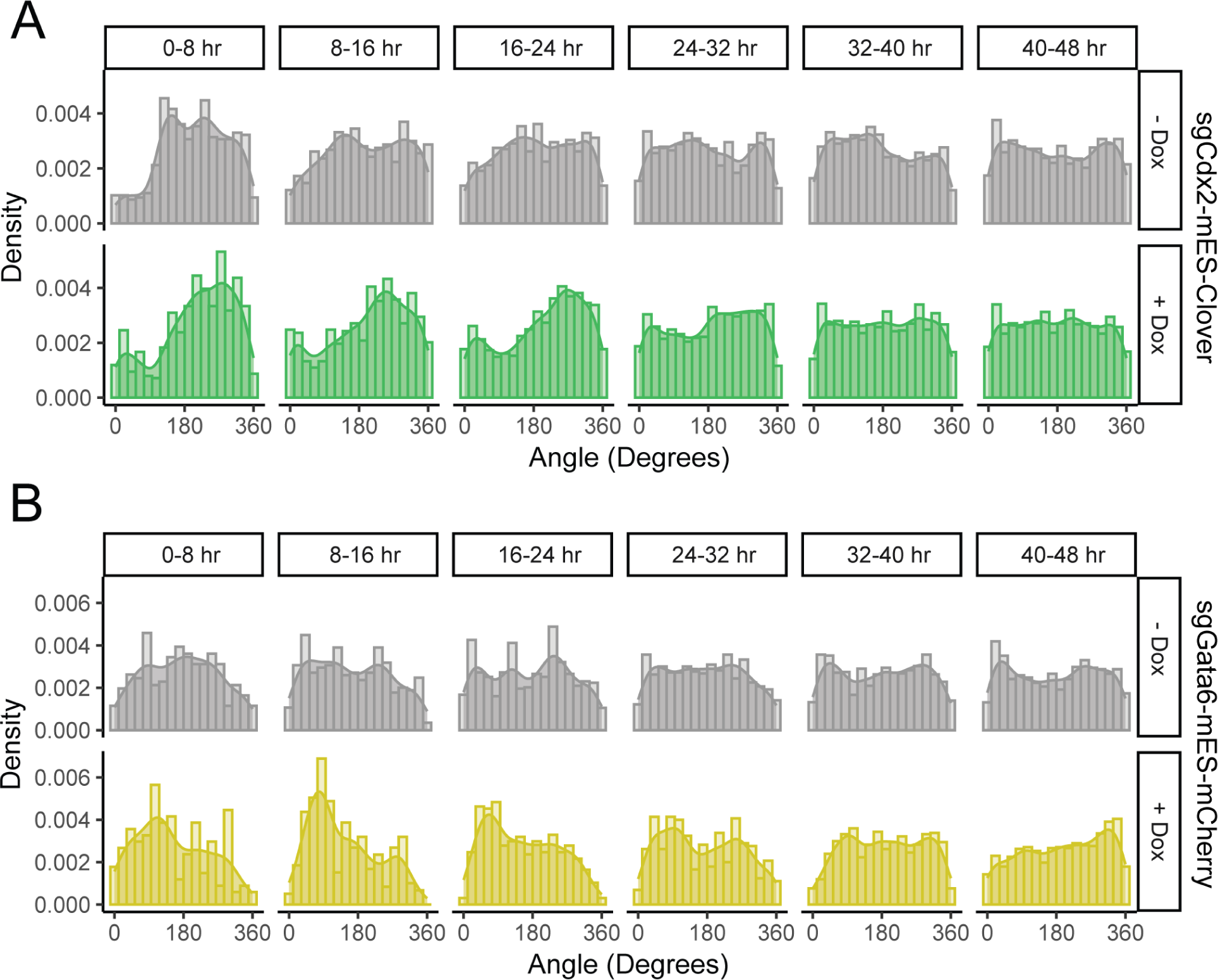
Additional plots on angle movement of cells. **A)** Distribution of movement angles for sgCdx2-mESCs, control (top) or doxycycline treated (bottom). **B)** Distribution of movement angles for sgGata6-mESCs, control (top) or doxycycline treated (bottom).

**Supplemental Figure 4.**
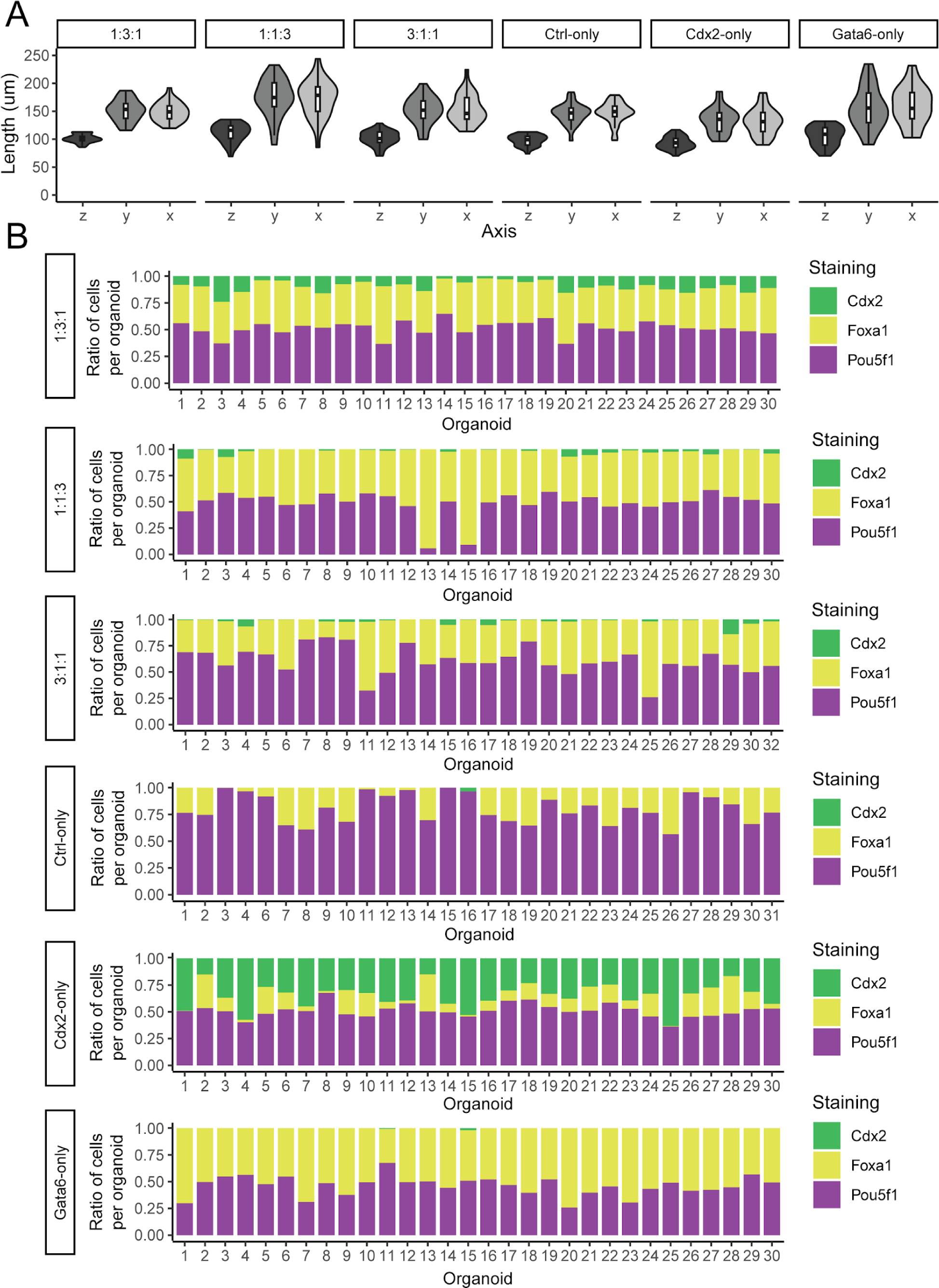
CPEM size distribution and relative cell distribution using different growth ratios (sgCtrl:sgCdx2:sgGata6 mESCs) and single cell-type only CPEMs. **A)** Size in micrometers of CPEMs in x-, y- and z-axis, longest axis within CPEMs for each, across all CPEM growth conditions analyzed as indicated. n=30 for each condition**. B)** Ratio of cells labeled Cdx2+, Foxa1+ and Pou5f1+ per region, across all CPEM growth conditions analyzed as indicated, n=30 for each condition.

**Supplemental Figure 5.**
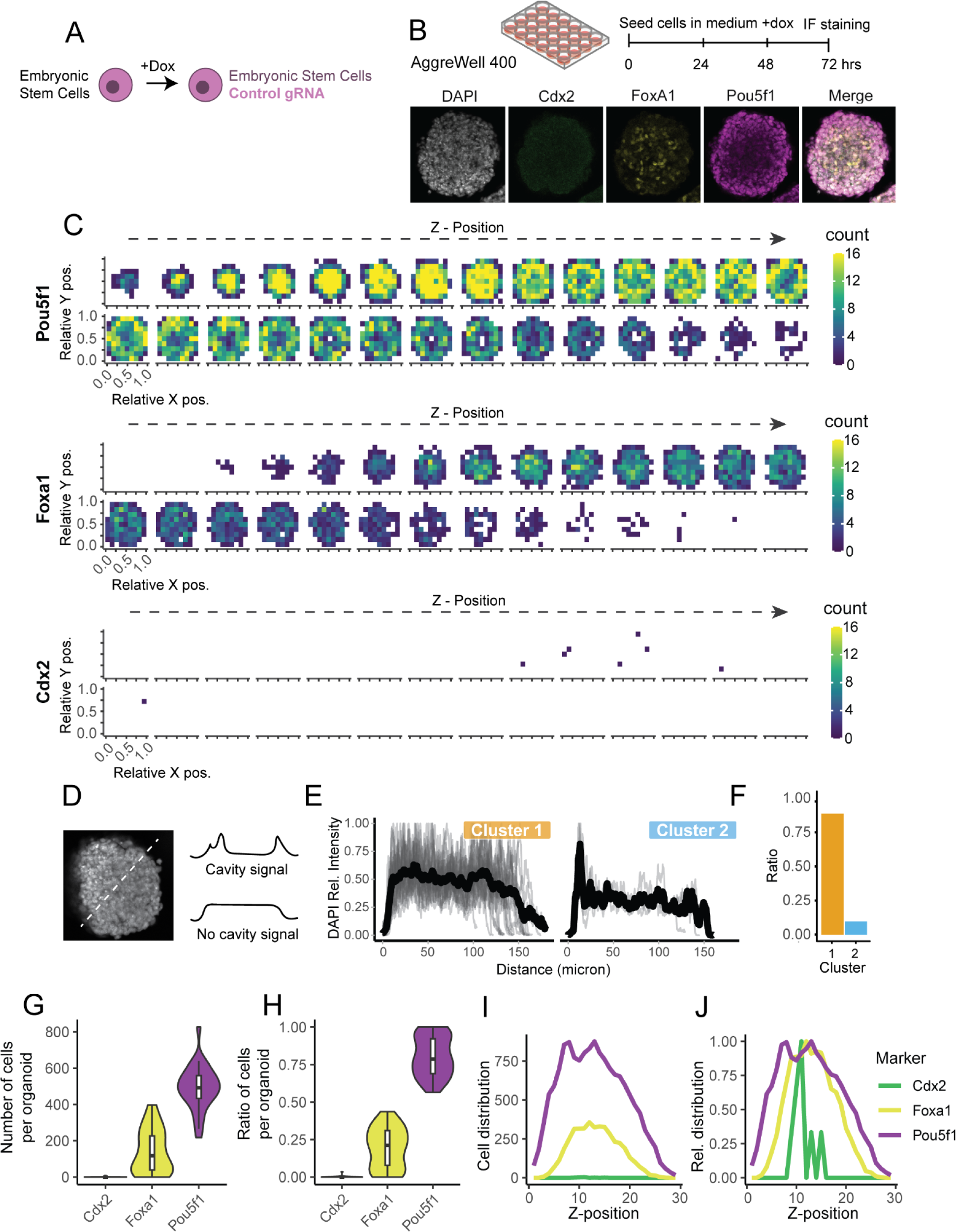
Generation and single-cell analysis of CRISPRa-programmed embryo models from sgCtrl-mESCs only. **A)** Overview of cells used for differentiation **B)** Differentiation protocol and representative confocal image of a 3D CPEMs with each cell type present, stained by Pou5f1, Cdx2 and Foxa1. **C)** Quantification of cell density for each marker across CPEMs using z-stack confocal imaging (n=30, n=19,017 nuclei labels), left to right shows increasing depth with 3.89um z-stack step-size, 28 density maps across the z-axis are displayed. **D)** Analysis of cavity formation by analyzing DAPI stain in the center position stack of CPEMs. **E)** Quantification and clustering of cavity formation in 2 clusters, and **F)** ratio of the clusters (n=30). **G)** Number and **H)** ratio of Cdx2, Foxa1 and Pou5f1 labeled cells identified in CPEMs (n=30). **I)** Mean number of Cdx2, Foxa1 and Pou5f1 labeled cells across imaged z-positions (n=30). **J)** Relative distribution of Cdx2, Foxa1 and Pou5f1 labeled cells across imaged z-positions (n=30), scaled to maximum within each group.

**Supplemental Figure 6.**
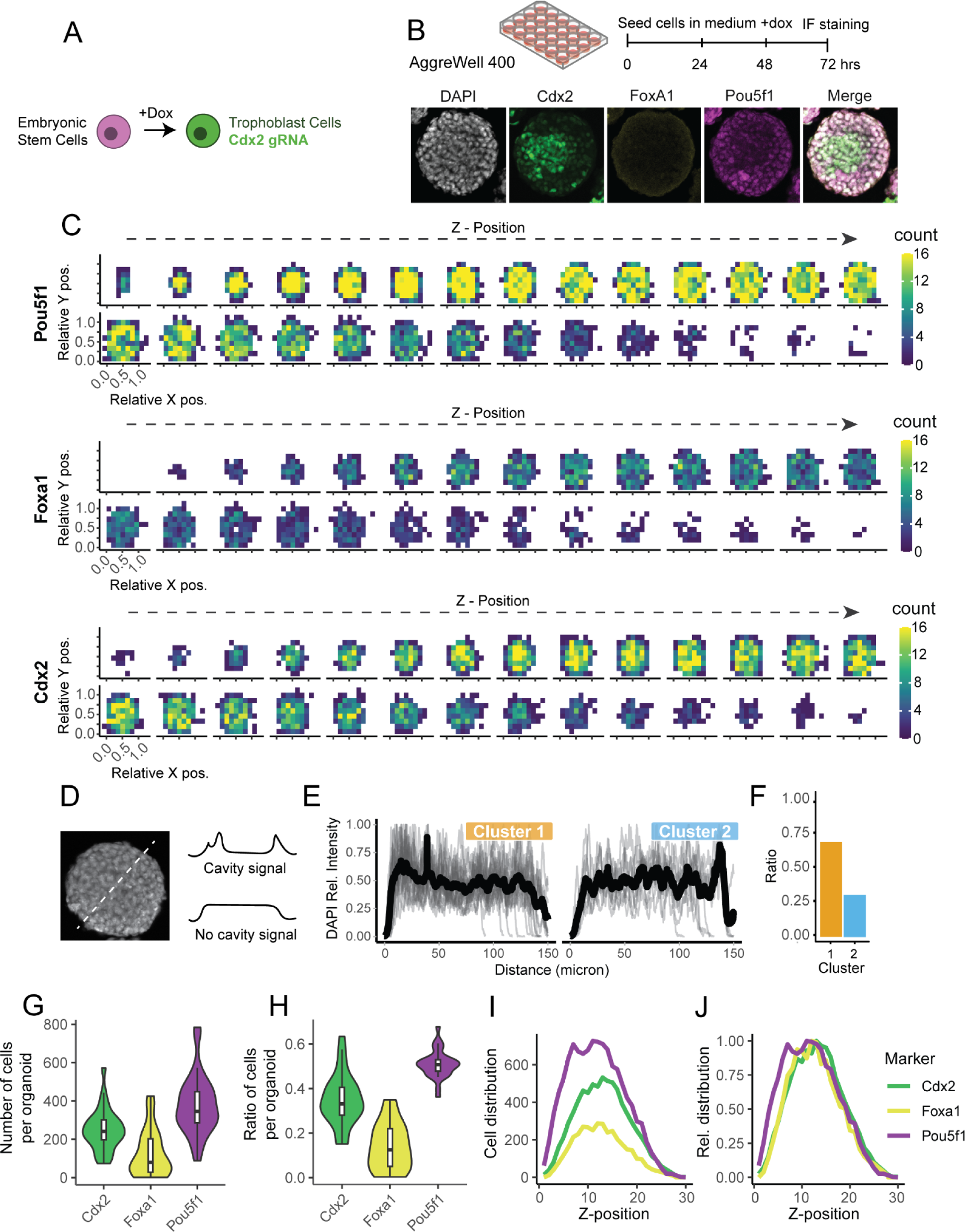
Generation and single-cell analysis of CRISPRa-programmed embryo models from sgCdx2-mESCs only. **A)** Overview of cells used for differentiation **B)** Differentiation protocol and representative confocal image of a 3D CPEM with each cell type present, stained by Pou5f1, Cdx2 and Foxa1. **C)** Quantification of cell density for each marker across CPEMs using z-stack confocal imaging (n=30, n=22,559 nuclei labels), left to right shows increasing depth with 3.89um z-stack step-size, 28 density maps across the z-axis are displayed. **D)** Analysis of cavity formation by analyzing DAPI staining in the center position stack of CPEMs. **E)** Quantification and clustering of cavity formation in 2 clusters, and **F)** ratio of the clusters (n=30). **G)** Number and **H)** ratio of Cdx2, Foxa1 and Pou5f1 labeled cells identified in CPEMs (n=30). **I)** Mean number of Cdx2, Foxa1 and Pou5f1 labeled cells across imaged z-positions (n=30). **J)** Relative distribution of Cdx2, Foxa1 and Pou5f1 labeled cells across imaged z-positions (n=30), scaled to maximum within each group.

**Supplemental Figure 7.**
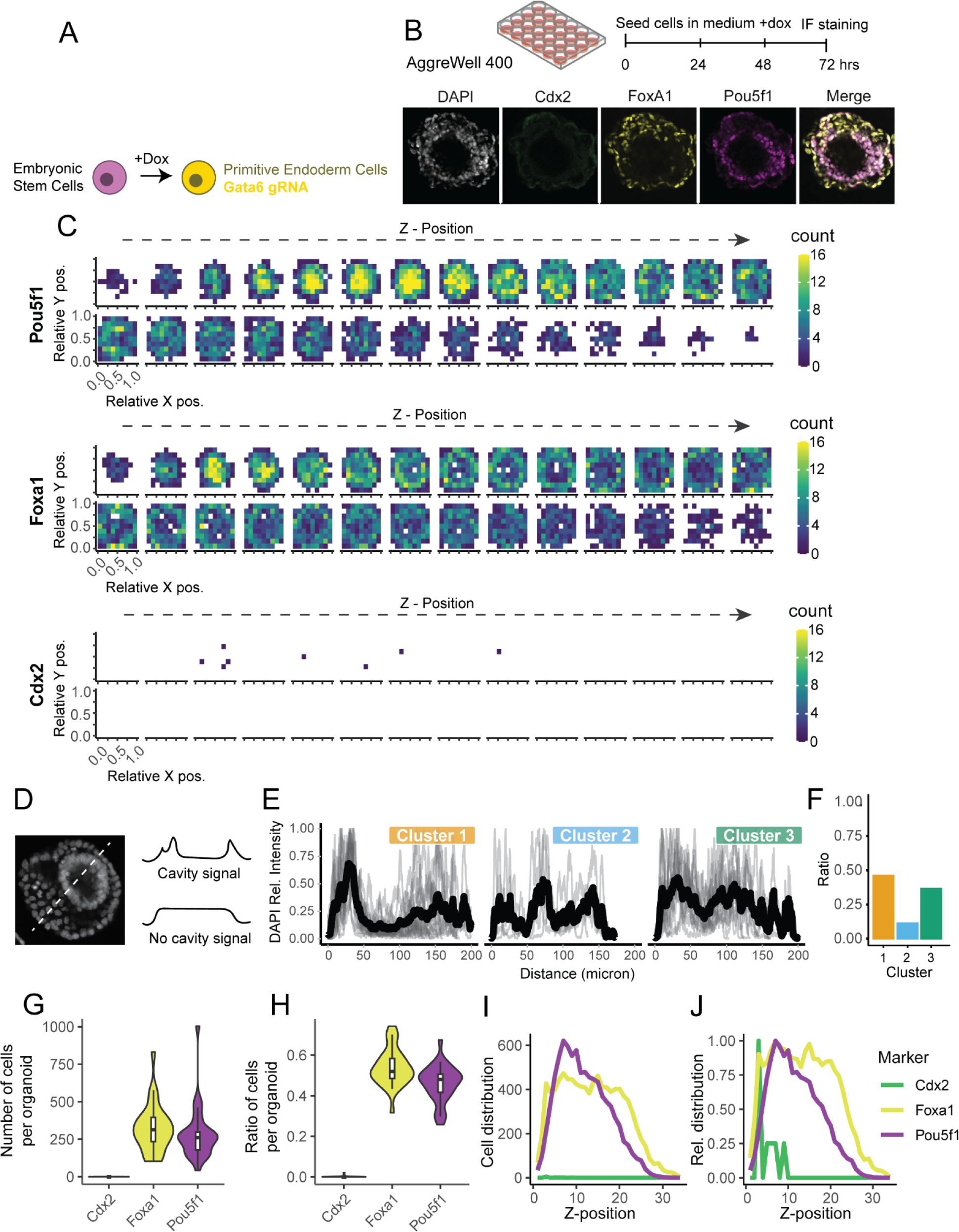
Generation and single-cell analysis of CPEMs from sgGata6-mESCs only. **A)** Overview of cells used for differentiation **B)** Differentiation protocol and representative confocal image of a 3D CPEM with each cell type present, stained by Pou5f1, Cdx2 and Foxa1. **C)** Quantification of cell density for each marker across CPEMs using z-stack confocal imaging (n=30, n=18,245 nuclei labels), left to right shows increasing depth with 3.89um z-stack step-size, 28 density maps across the z-axis are displayed. **D)** Analysis of cavity formation by analyzing DAPI stain in the center position stack of CPEMs. **E)** Quantification and clustering of cavity formation in 2 clusters, and **F)** ratio of the clusters (n=30). **G)** Number and **H)** ratio of Cdx2, Foxa1 and Pou5f1 labeled cells identified in CPEMs (n=30). **I)** Mean number of Cdx2, Foxa1 and Pou5f1 labeled cells across imaged z-positions (n=30). **J)** Relative distribution of Cdx2, Foxa1 and Pou5f1 labeled cells across imaged z-positions (n=30), scaled to maximum within each group.

**Supplemental Figure 8.**
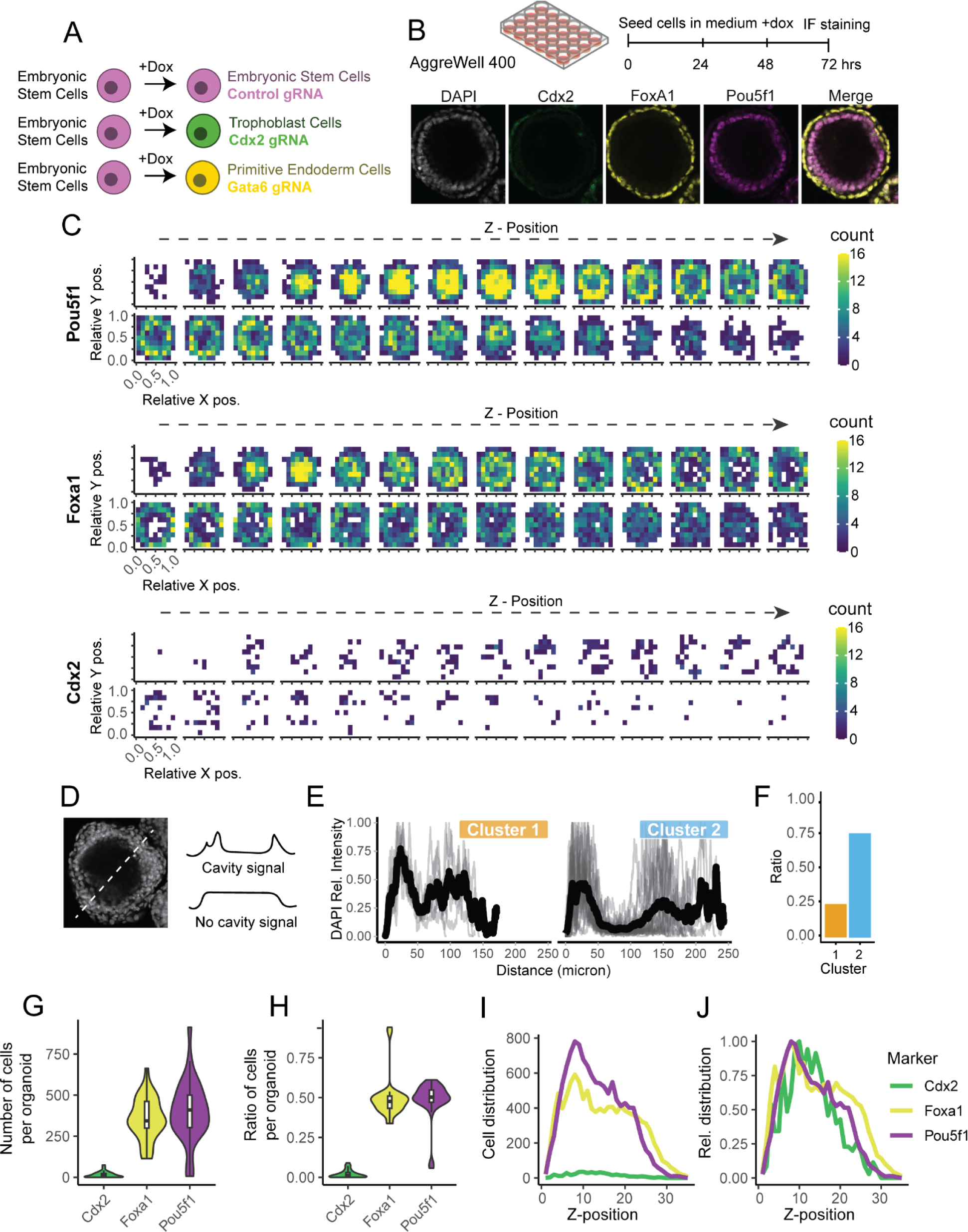
Generation and single-cell analysis of CRISPRa-programmed embryo models from a 1:1:3 ratio of cells (sgCtrl:sgCdx2:sgGata6 mESCs). **A)** Overview of cells used for differentiation **B)** Differentiation protocol and representative confocal image of a CPEM with each cell type present, stained by Pou5f1, Cdx2 and Foxa1. **C)** Quantification of cell density for each marker across CPEMs using z-stack confocal imaging (n=30 CPEMs, n=23,156 nuclei labels), left to right shows increasing depth with 3.89um z-stack step-size, 28 density maps across the z-axis are displayed. **D)** Analysis of cavity formation by analyzing DAPI stain in the center position stack of CPEMs. **E)** Quantification and clustering of cavity formation in 2 clusters, and **F)** ratio of the clusters (n=30). **G)** Number and **H)** ratio of Cdx2, Foxa1 and Pou5f1 labeled cells identified in CPEMs (n=30). **I)** Mean number of Cdx2, Foxa1 and Pou5f1 labeled cells across imaged z-positions (n=30). **J)** Relative distribution of Cdx2, Foxa1 and Pou5f1 labeled cells across imaged z-positions (n=30), scaled to maximum within each group.

**Supplemental Figure 9.**
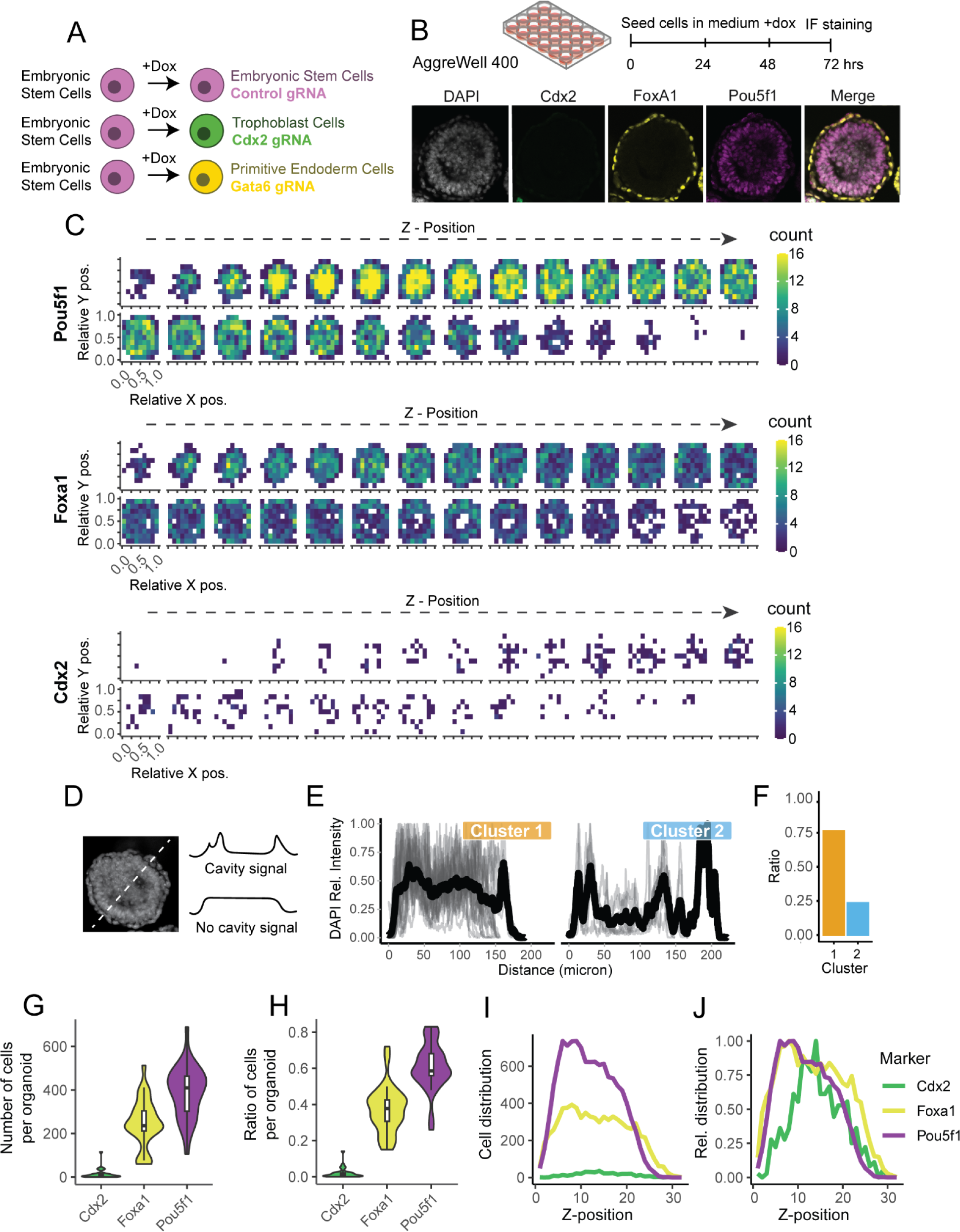
Generation and single-cell analysis of CRISPRa-programmed embryo models from a 3:1:1 ratio of cells (sgCtrl:sgCdx2:sgGata6 mESCs). **A)** Overview of cells used for differentiation **B)** Differentiation protocol and representative confocal image of a CPEM with each cell type present, stained by Pou5f1, Cdx2 and Foxa1. **C)** Quantification of cell density for each marker across CPEMs using z-stack confocal imaging (n=30, n=19,538 nuclei labels), left to right shows increasing depth with 3.89um z-stack step-size, 28 density maps across the z-axis are displayed. **D)** Analysis of cavity formation by analyzing DAPI stain in the center position stack of CPEMs. **E)** Quantification and clustering of cavity formation in 2 clusters, and **F)** ratio of the clusters (n=30). **G)** Number and **H)** ratio of Cdx2, Foxa1 and Pou5f1 labeled cells identified in CPEMs (n=30). **I)** Mean number of Cdx2, Foxa1 and Pou5f1 labeled cells across imaged z-positions (n=30). **J)** Relative distribution of Cdx2, Foxa1 and Pou5f1 labeled cells across imaged z-positions (n=30), scaled to maximum within each group.30), scaled to maximum within each population.

**Supplemental Figure 10.**
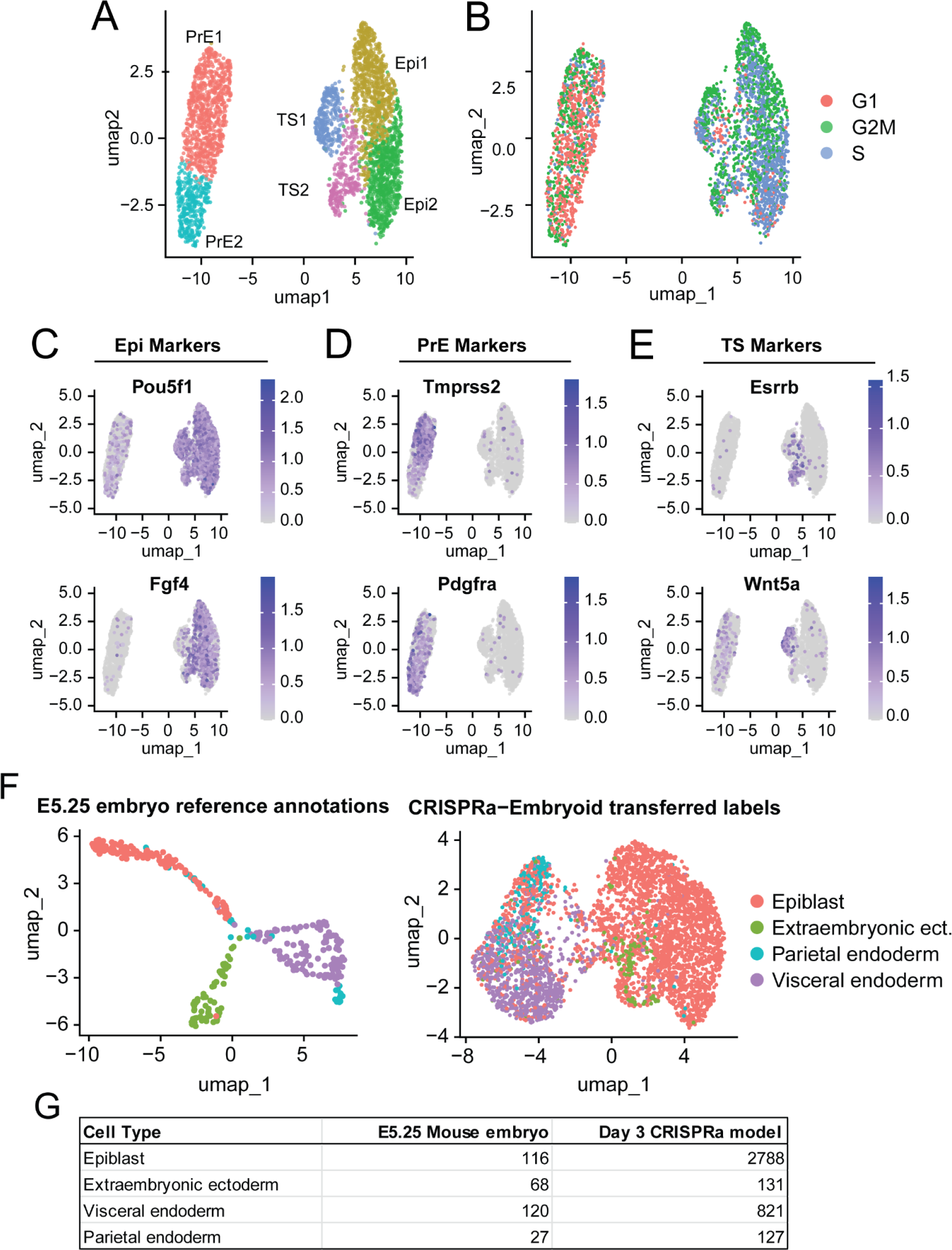
Additional features of scRNA-seq data of mixed cell line CPEMs and comparison with *in vivo* mouse embryo data. **A)** Umap of cell clusters identified in mixed cell line CPEMs. **B)** Feature plot showing predicted cell cycle phases in mixed cell 3D CPEMs. **C)** Feature plots of additional Epiblast markers. **D)** Feature plots of additional Primitive Endoderm markers. **E)** Feature plots of additional Trophoblast markers. **F)** Umap of labels transferred between E5.25 mouse embryo data (left) and mixed cell embryoid data (right). **G)** Data table showing mapping of cell labels between datasets.

**Supplemental Figure 11.**
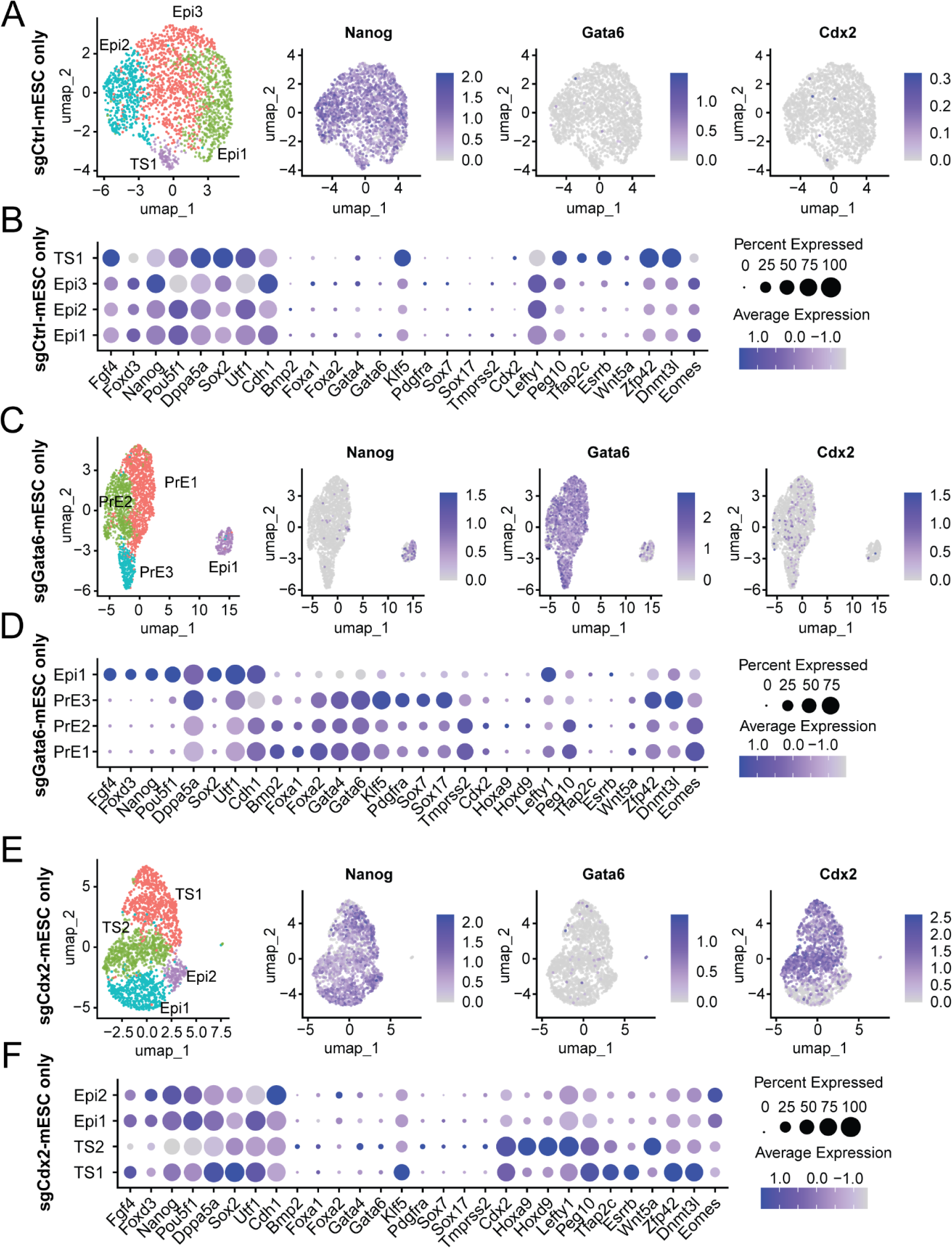
Summary of scRNA-seq data of CPEMs grown from individual cell lines. **A)** Umap and featureplots of CPEMs grown from sgCtrl-mESCs only. **B)** Umap and featureplots of CPEMs grown from sgGata6-mESCs only. **C)** Umap and featureplots of 3D CPEMs grown from sgCdx2-mESCs only. **D)** Dotplot for marker genes of Epiblast, Primitive Endoderm and Trophoblast cells in sgCtrl-mESC only CPEM data. **E)** Dotplot for marker genes of Epiblast, Primitive Endoderm and Trophoblast cells in sgGata6-mESC only CPEM data. **F)** Dotplot for marker genes of Epiblast, Primitive Endoderm and Trophoblast cells in sgCdx2-mESC only CPEM data.

**Supplemental Figure 12.**
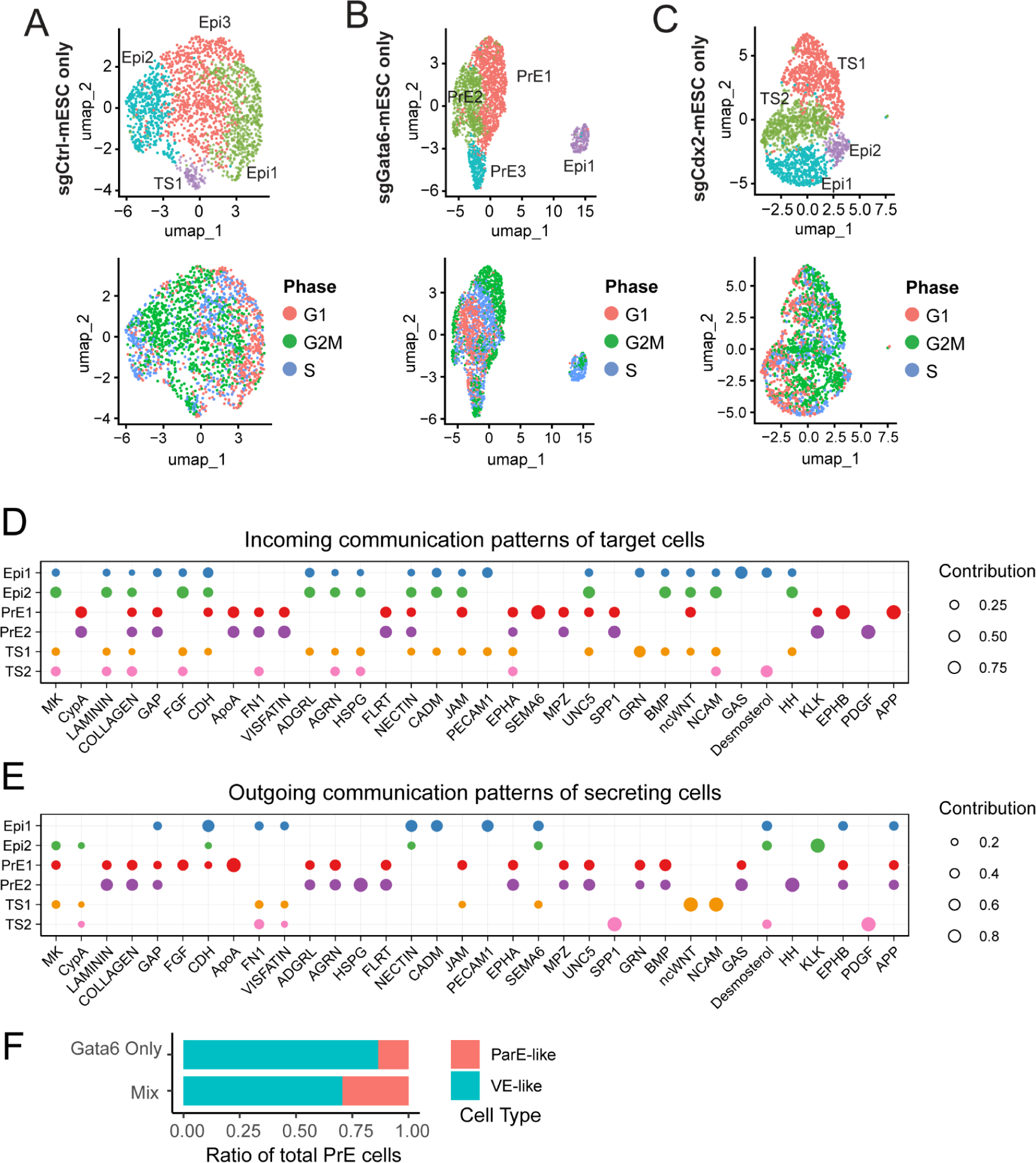
Cell cycle analysis of single cell type CPEMs and CellChat mixed CPEMs pathway activity profiles. **A)** Cell cycle profiles in sgCtrl-mESC only CPEMs. **B)** Cell cycle profiles in sgGata6-mESC only CPEMs. **C)** Cell cycle profiles in sgCdx2-mESC only CPEMs. **D)** Incoming signaling per cell cluster, all observed interactions from CellChat. **E)** Outgoing signaling per cluster, all observed interactions from CellChat. **F)** Comparison of the ratio of parietal-like and visceral-like endoderm cells in mixed CPEMs and sgGata6-mESC only CPEMs.

**Supplemental Movie 1.**

**A)** Complete time lapse imaging for live cell data shown in Figure 3B. Cell tracking data (Btrack) is overlaid on fluorescent signals and treatments as indicated in movie. **B)** Additional time lapse imaging examples for live cell data shown in Figure 3, showing 5 individual micropattern regions for doxycycline and control treated wells. Cell segmentation (stardist) and tracking data (Btrack) is overlaid on fluorescent signals and treatments as indicated.

**Supplemental Movie 2.**

**A)** Example of Z-projection of a CPEM from mixed cells (1:3:1, sgCtrl:sgCdx2:sgGata6 mESCs). **B)** Example of Z-projection of a CPEM from sgCtrl-mESCs only, stained for Pou5f1 (Magenta), Cdx2 (Cyan) and Foxa1 (Yellow). **C)** Example of Z-projection of a CPEM from sgCdx2-mESCs only, stained for Pou5f1 (Magenta), Cdx2 (Cyan) and Foxa1 (Yellow). **D)** Example of Z-projection of a CPEM from sgGata6-mESCs only, stained for Pou5f1 (Magenta), Cdx2 (Cyan) and Foxa1 (Yellow). For each Z-projection, half of the images in the z-axis are shown to display both inside and outside structures of CPEM structures.

## Methods and Materials

### Cell culture

Mouse embryonic stem cells (mESCs) were cultured in 2i+Lif conditions in either ESGRO-2i medium (MilliporeSigma, SF016-200) +1x Pen/Strep (Thermo Fisher Scientific, 15140122) or N2B27 medium (Takara, Y40002) supplemented with 1x Pen/Strep (Thermo Fisher Scientific, 15140122), 1uM PD0325901 (StemRD), 3uM CHIR99021 (StemRD) and 1000 U/ml Lif (MilliporeSigma, ESG1107). mESCs were grown on tissue culture treated plates coated with 0.1% gelatin (Sigma G1890). For routine passaging, cells were seeded at 5,000-10,000 cells per cm^2^ and passaged every 2-3 days using Accutase (Innovative Cell Technologies, AT104).

### Generation of an inducible CRISPRa cassette

For integrating the PB-TRE3G-dCas9-VPR cassette (dCas9-VPR) (Addgene, #63800), cells were transfected using Lipofectamine Stem (Thermo Fisher Scientific, STEM00003), diluted in OptiMEM (Thermo Fisher, 31985062). Briefly, mESCs were seeded in 0.1% gelatin coated 6 well plates as described before, seeding 300,000 cells per well. The next day, cells were transfected with the following plasmids diluted in OptiMEM : PiggyBac Transposase (200 ng), dCas-VPR (800 ng) and pUC19 (1000 ng). As a non-integrating control transfection, the following condition was used: dCas-VPR (800 ng) and pUC19 (1200 ng). Medium was replaced 6 hours after transfection. 48 hours after transfection, cells with successful integration were selected by addition of 100 ug/ml Hygromycin (Mirus Bio, MIR5930) for 4 days, or until no control cells were remaining.

### Lentiviral transduction of sgRNA cassettes

The pSLQ1371-sgRNA cassette (Addgene, 121425) was integrated via lentiviral transduction. To prepare lentivirus, HEK293T cells were seeded on 6 well plates at a density of 500,000 cells per well, in medium consisting of DMEM+Glutamax (Thermo Fisher Scientific 10566024), 1x Pen/Strep (Thermo Fisher Scientific, 15140122) and 10% HIFBS (Thermo Fisher Scientific, 10438026). The next day, per well, medium was replaced to 2ml DMEM+Glutamax (Thermo Fisher Scientific 10566024) and 10% ESC qualified FBS (MilliporeSigma, ES-009-B). Cells were then transfected using Turbofect (Thermo Fisher Scientific,) with the following plasmids: VSVG (300 ng), DR8.74 (450 ng), pSLQ1371-sgRNA (750 ng). 6 hours after transfection, 1 ml DMEM+Glutamax (Thermo Fisher Scientific 10566024), 3x Pen/Strep (final concentration 1x) (Thermo Fisher Scientific, 15140122), and 10% ESC qualified FBS (MilliporeSigma, ES-009-B) was added. Lentivirus was harvested 72 hours after transfection by collecting the medium into tubes, followed by centrifugation at 500 rcf. Next, the supernatant was filtered through a 0.45 um filter (MilliporeSigma, SLHVM33RS) and collected into new tubes.

For lentiviral transduction, mESCs were seeded on 0.1% gelatin coated 6 well plates at a density of 50,000 cells per well. The next day, medium was replaced with 1 ml 2i+Lif medium and 1 ml prepared lentivirus. After 24 hours of incubating the lentivirus, medium was replaced again to 2i+Lif only medium. To select for transduced cells, 0.8 ug/ml Puromycin (Stemcell Technologies, 73342) was added 48 hours after lentiviral transduction, and added for 4 days or until no control cells (non-transduced cells) were remaining.

Using the above methods, the 3 mESC lines were generated with a control non-targeting gRNA, Gata6 targeting gRNA and Cdx2 targeting gRNA [80] (**Supplemental Table 1, Supplemental Figure 1A**), named sgCtrl-mESC, sgGata6-mESC and sgCdx2-mESC respectively.

**Supplemental Table 1.**
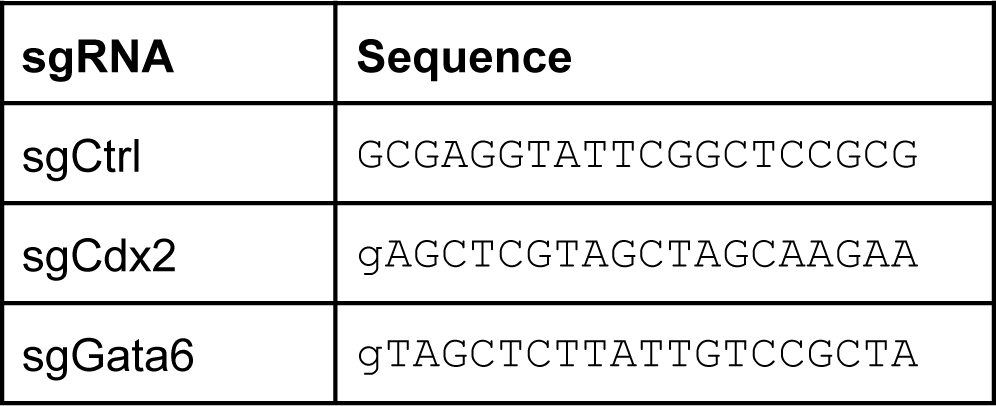
gRNAs used in sgCtrl-mESCs, sgCdx2-mESCs and sgGata6-mESCs.

### CRISPRa validation immunofluorescence staining

For validation of the 3 CRISPRa lines, induction of target genes was done by immunofluorescence staining. mESCs were seeded 0.1% gelatin coated 24 well plates. Depending on experiment duration, cells were seeded at the following densities: 24 hour treatment - 50,000 cells per well, 48 hour treatment - 25,000 cells per well, 72 hour treatment - 12,500 cells per well, 96 hour treatment - 6,250 cells per well. For each staining sgCtrl-mESC, sgGata6-mESC and sgCdx2-mESC were seeded in duplicate to perform doxycycline and no doxycycline control treatments. Cells were seeded in standard differentiation medium: DMEM+Glutamax, 15% ESC qualified FBS, 1x Sodium Pyruvate (Thermo Fisher, 11360070), 1x Non-essential amino acids (Thermo Fisher, 11140050), 1x Pen/Strep and 0.1 mM beta-mercaptoethanol (Thermo Fisher, 21985023). Doxycycline was added at 0.5 ug/ml (Millipore Sigma, D5207) in treatment wells. Medium was changed every 24 hours until ready for immunofluorescence staining. Cells were prepared by adding 0.4 ml of 3.6% Formaldehyde (Avantor, JTS898-7) in 1x PBS and incubating for 10 minutes at room temperature. Each well was washed twice with 1x PBS and incubated with 0.2% Triton-X100 (VWR, 97063-864) in 1x PBS for 20 minutes. Each well was washed once with 1x PBS and incubated with 0.4 ml Blocking buffer consisting of 3% BSA + 0.1% Triton in 1x PBS (Blocking buffer) for 1 hour. Blocking buffer was removed, 0.4 ml of Primary antibody incubation solution was added per well, consisting of 1% BSA in 1x PBS, including diluted antibodies according to the table below. Plates were incubated overnight at 4°C.

| **Antibody** | **Cat. No.** | **Dilution** |
| --- | --- | --- |
| Rabbit anti Cdx2 | Cell Signaling Technology, 12306S | 1:2000 |
| Rabbit anti Gata6 | Cell Signaling Technology, 5851S | 1:2000 |
| Rabbit anti Gata3 | Cell Signaling Technology, 5852S | 1:1000 |
| Goat anti Tfap2c | R&D Systems, AF5059 | 1:1000 |
| Mouse anti Foxa1 | Abcam, ab55178 | 1:2000 |
| Goat anti Sox17 | R&D Systems, AF1924 | 1:2000 |
| Goat anti Oct3/4 | R&D Systems, AF1759 | 1:1000 |
| Mouse anti Oct3/4 | Santa Cruz Biotechnology, sc-5279 | 1:200 |

The next day, Primary antibody incubation solution was removed and wells were washed twice with 0.1% Tween in 1x PBS and washed once with 1x PBS. 0.4 ml of Secondary antibody solution was used, consisting of 1% BSA in 1x PBS, including diluted antibodies according to the table below. Plates were incubated at room temperature for 1 hour. Secondary antibody solution was removed and wells were incubated with 0.5 ug/ml DAPI + 0.1% Tween (BioRad,1610781) in 1x PBS for 5 minutes. Wells were washed once with 0.1% Tween in 1x PBS and twice with 1x PBS. For imaging, 0.5 ml of 1x PBS was added per well. All washing steps were 3 minutes each and approximately 0.5 ml volume unless otherwise indicated. Immunofluorescence and phase contrast images were captured using the Lionheart FX system (Agilent BioTek) with Gen5 software (3.14). 16 images were captured per well, using phase contrast, DAPI, GFP and TexasRed channels. Images were processed in imageJ. Image stacks were loaded per channel and a binary mask was created using DAPI stain for each condition, to identify individual nuclei regions. Then, the DAPI mask was used to measure intensity of stainings in GFP and TexasRed channels. The mean intensity per nucleus for each staining was extracted in this way for all conditions and plotted using the ggplot2 [81] and ggridges packages in Rstudio.

| Antibody | Cat. No. | Dilution |
| --- | --- | --- |
| Donkey anti Rabbit Alexa 488 | Thermo Fisher, A21206 | 1:1000 |
| Donkey anti Goat Alexa 488 | Thermo Fisher, A11055 | 1:1000 |
| Donkey anti Rabbit Alexa 594 | Thermo Fisher, A21207 | 1:1000 |
| Goat anti Mouse Alexa 488 | Thermo Fisher, A11001 | 1:1000 |
| Goat anti Mouse Alexa 594 | Thermo Fisher, A11005 | 1:1000 |

### Micropattern culture and immunofluorescence staining

For micropattern culture and end-point immunofluorescence staining, Cytoo Arena A microchips (CYTOO, 10-020-00-18) were used. Chips were placed into 6 well plates already containing 3 ml of 1x PBS. Then, 1x PBS was removed and 3 ml of 4 ug/ml Laminin in 1x PBS was added and placed in a cell culture incubator for 2 hours at 37°C, 5% CO2. Wells were washed by adding 6 ml 1x PBS per well and removing 6 ml solution, repeated 5 times. Then, one well at a time, complete solution was removed and 3 ml 2i+Lif medium + 10uM ROCK inhibitor (Stemcell Technologies, 72304) was added. Plates were placed in the incubator until needed the same day. To plate cells, sgCtrl-mESCs, sgGata6-mESCs and sgCdx2-mESCs were individually collected, counted, and mixed together in a 1:1:1 ratio. Then, 120,000 cells were added per well (40,000 for each line). After 2 hours, medium was replaced to standard differentiation medium: DMEM+Glutamax, 15% ESC qualified FBS, 1x Sodium Pyruvate, 1x Non-essential amino acids, 1x Pen/Strep and 0.1 mM beta-mercaptoethanol, and doxycycline was added to treatment wells. Medium was changed daily, one well at a time. Cells were grown on microchips for 72 hours before immunofluorescence staining.

To prepare Cytoo Arena A microchips for immunostaining, medium was removed and wells were washed once with 1x PBS, followed by addition of 3 ml 3.6% formaldehyde in 1x PBS per well and incubated 30 minutes at room temperature. Each well was washed twice with 1x PBS and microchips were transferred from 6 well plates into individual 35mm plates per microchip, already containing 3 ml 1x PBS. The solution was replaced with 3 ml 0.2% Triton in 1x PBS and incubated for 30 minutes at room temperature. Each dish was washed once with 1x PBS and incubated with 3 ml Blocking buffer consisting of 3% BSA + 0.1% Triton in 1x PBS (Blocking buffer) for 2 hours. Blocking buffer was removed, 2.5 ml of Primary antibody incubation solution was added per dish, consisting of 1% BSA in 1x PBS, including diluted antibodies according to the table below. Dishes were incubated overnight at 4°C.

| Antibody | Cat. No. | Dilution |
| --- | --- | --- |
| Rabbit anti Cdx2 | Cell Signaling Technology, 12306S | 1:2000 |
| Mouse anti Foxa1 | Abcam, ab55178 | 1:2000 |
| Goat anti Oct3/4 | R&D Systems, AF1759 | 1:1000 |

The next day, Primary antibody incubation solution was removed and dishes were washed twice with 0.1% Tween in 1x PBS and washed once with 1x PBS. Dishes were incubated for 2 hours at room temperature in 2.5 ml of Secondary antibody solution, consisting of 1% BSA in 1x PBS, including diluted antibodies according to the table below. Secondary antibody solution was removed and dishes were incubated with 0.5 ug/ml DAPI + 0.1% Tween in 1x PBS for 10 minutes. Dishes were washed once with 0.1% Tween in 1x PBS and twice with 1x PBS. For imaging, 3 ml of 1x PBS was added per dish. All washing steps were 5 minutes each and 1 ml volume unless otherwise indicated. Immunofluorescence images were captured using the Zeiss LSM<880 confocal laser scanning microscope system.

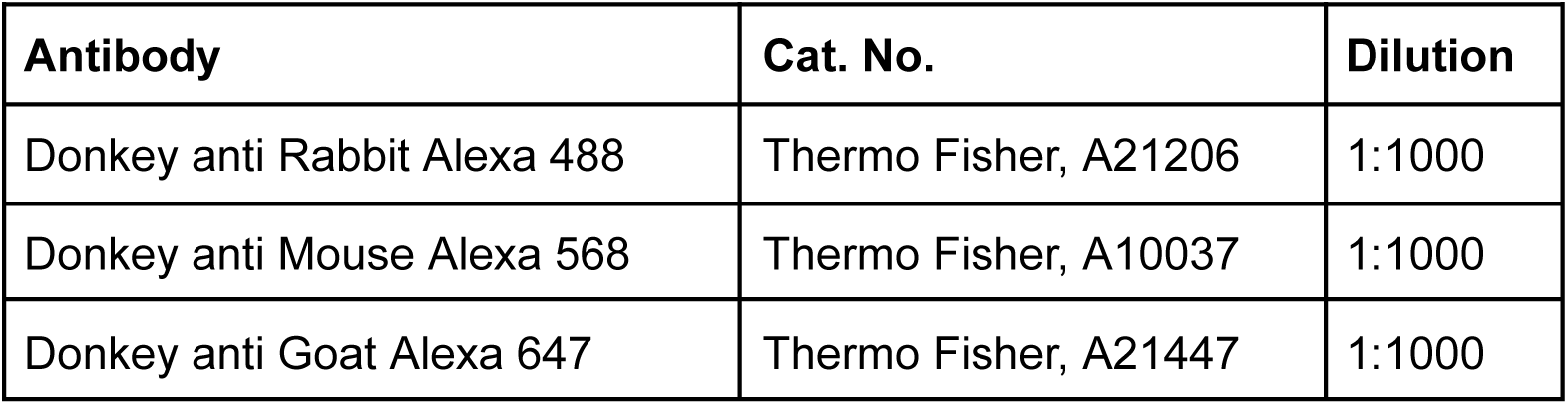

### Micropattern image segmentation and processing

For each channel, tiled image scans were stitched, followed by maximum projection intensity of z-stacks (11 image stack, 3.19um z-intervals), in Zeiss ZEN software. The resulting images were imported into Napari per channel and segmented using StarDist (stardist-napari 2022.12.6). The following settings were used: Model Type: 2D. Pre-trained model: Versatile (fluorescent nuclei). Normalize image yes, percentile low = 1.00, percentile high = 99.80. Probability/Score threshold: 0.6. Overlap threshold: 0.5. Output type: Labels only. Normalization axes: ZYX. Time-lapse labels: Match to previous frame (via overlap). Individual regions were extracted and cropped using square shape layers for each region. For each channel, the segmentation data was analyzed with RegionProps (napari-skimage-regionprops 0.10.1) to extract positions, size and signal intensity of each individually segmented cell and exported to a data table. The number and ratio of cells for each region was plotted using ggplot2. Density plots were made using the ggplot2 geom_bin2d option, using 15 bins. For each region, using centroid data, the x/y coordinates were scaled between 0-1, where 0 was the minimum pixel position where the first cell was along the x/y axis, and 1 the maximum position where the last cell was along the x/y axis, to account for small variations in size and crops, to better capture the average pattern.

### Live cell micropattern imaging

For live cell imaging and tracking of different co-cultured mESC lines, the pSLQ1371 sgRNA plasmid was modified by replacing mCherry for either Clover or mTagBFP. The Myc nuclear localization signal (NLS) was also added to the C-terminus each fluorescent protein (mCherry-NLS, Clover-NLS and mTagBFP-NLS). The mESC line with stably integrated inducible dCas9-VPR was transduced via lentivirus with DNA encoding for either sgCtrl-mTagBFP-NLS, sgGata6-mCherry-NLS, or sgCtrl-Clover-NLS, as described before.

For live cell micropattern culture, RosetteArray™ (Neurosetta, Spinal 6-Well 100µm RosetteArray™ Plate) 6 well plates were used with 100 um growth regions. Wells were washed once with 1x PBS, followed by incubating 3 ml of 4 ug/ml Laminin in 1x PBS for 6 hours in a cell culture incubator for 6 hours at 37°C, 5% CO2. Wells were washed once with 3 ml 1x PBS per well one well at a time. Then, one well at a time, complete solution was removed and 2 ml standard differentiation medium was added. Plates were placed in the incubator until needed the same day. sgCtrl-mTagBFP-NLS-mESCs, sgGata6-mCherry-NLS-mESCs and sgCdx2-Clover-NLS-mESCs were individually collected, counted, and mixed together in a 1:1:1 ratio. Then, 400,000 cells were added per well (133,000 for each line) in a volume of 1 ml, so the total volume per well was 3 ml. After 6 hours, doxycycline was added to treatment wells. Medium was changed daily, one well at a time. For live cell imaging, the plate was placed in a Lionheart FX imager equipped with a humidity chamber, temperature (37°C) and CO2 (5%) control. Cells were imaged every hour for 48 hours total, using phase contrast, YFP and TexasRed channels.

### Live cell micropattern image segmentation and processing

Tiled images obtained by Lionheart FX were stitched using Gen5 3.14 software. Time-lapse data was imported into imageJ as image stacks and individual micropattern regions were cropped and saved as individual image sequences per channel. For each micropattern region, the time-lapse data was imported as stacks into Napari. Cells were segmented using StarDist with label over time option. Next, Btrack plugin (0.6.4) was used to track individual cells in timelapse data and exported as h5 files. Default Btrack “cell” settings were used, except Motion Accuracy was increased to 10. For videos of live cell data, layers were arranged and/or overlaid in Napari and exported using wizard plugin (napari-animation 0.0.7). For detailed analysis of cell dynamics, cell displacement data was extracted from h5 files into .csv file format and further processed using CelltrackR (1.1.0)[82] to calculate mean square displacement, and angles were calculated in reference to the center of each micropattern region between individual cell data points in the timelapse imaging data.

### CRISPRa-programmed embryo model culture and immunofluorescence staining

To grow and differentiate mESCs in 3D structures, cells were grown in 24 well AggreWell 400 plates (Stemcell Technologies, 34415). Each well was washed once with 1x PBS, followed by adding 0.5 ml of anti-adherence solution (Stemcell Technologies, 07010). Plates were centrifuged 5 minutes at 1,000 rcf and incubated 20 minutes at room temperature. Anti-adherence solution was removed and 1 ml 1x PBS was added to each well. Plates were kept at room temperature until seeding cells. Just before seeding, 1x PBS was removed and 1 ml of standard differentiation medium + 20 uM ROCK inhibitor was added. To treatment wells, 1 ug/ml doxycycline was added. sgCtrl-mESCs, sgGata6-mESCs and sgCdx2-mESCs were individually collected, counted, and mixed together in a 1:1:3 ratio. Then, 32,000 cells were added per well (6,000 sgCtrl, 6,000 sgGata6 and 20,000 sgCdx2) in a volume of 1 ml, so that the total volume per well was 2 ml, and the final concentration of ROCKi was 10 uM and doxycycline was 0.5 ug/ml. After 24 hours, 1.4 ml of medium per well was replaced using standard differentiation medium without ROCKi, and repeated once. After another 24 hours 1.4 ml of medium was replaced per well. Doxycycline was added every medium change for treatment conditions. For conditions with altered cell ratios, the amount of cells was adjusted accordingly.

After 72 hours in culture, CMEMs were prepared for immunostaining. 1.5 ml of medium was removed per well, and 1.5 ml of 1x PBS was added. CPEMs were allowed to settle for 1 minute, then 1.5 ml of this solution was removed for each well. 0.5 ml of 1x PBS was added per well. By tapping the plate and/or pipetting up and down, using a cut-off P1000 tip to prevent shearing, CPEMs were suspended and then transferred to 6 well plates already containing 2 ml of 1x PBS. A 40 um cell strainer (Corning, 087711), with the side tab cut off, was added to each well, so it forms a barrier to prevent update of CPEMs when aspiring solutions from the well. For aspirating solutions, the plate was tilted slightly leading to approximately 0.5 to 1 ml of solution remaining to keep CPEMss in suspension. This way, 1x PBS was removed and each well was washed 2 ml of 3.6% formaldehyde in 1x PBS, followed by adding 2 ml of 3.6% formaldehyde in 1x PBS per well and incubated for 40 minutes at room temperature on an orbital shaker. For incubations, cell strainers were removed for improved mixing of CPEMs and buffers. During washes, cell strainers were briefly removed and reinserted after adding each wash to promote mixing of solutions. Formaldehyde solution was aspirated, then washed once with 1x PBS + 0.01% Tween + 1x Glycine (Cell Signaling Technology, #7005), and washed twice with 1x PBS + 0.01% Tween. Wells were washed once with 1x PBS + 0.2% Triton-X100, followed by adding 2 ml of 1x PBS + 0.2 % Triton X-100 and incubated for 30 minutes at room temperature on an orbital shaker. Solution was removed and washed once with Blocking buffer (1x PBS + 3% BSA + 0.1% Triton-X100), followed by addition of 2 ml Blocking buffer and incubated for 2 hours at room temperature on an orbital shaker. Solution was removed and washed once with Antibody incubation buffer (1x PBS + 3% BSA), followed by addition of 2 ml of Primary antibody incubation solution per dish, including diluted antibodies according to the table below. Dishes were incubated overnight at 4°C on an orbital shaker.

| **Antibody** | **Cat. No.** | **Dilution** |
| --- | --- | --- |
| Rabbit anti Cdx2 | Cell Signaling Technology, 12306S | 1:1000 |
| Mouse anti Foxa1 | Abcam, ab55178 | 1:1000 |
| Goat anti Oct3/4 | R&D Systems, AF1759 | 1:500 |

Primary antibody solution was removed and wells were washed three times with 1x PBS + 0.1% Tween. Wells were washed once with an antibody incubation buffer (1x PBS + 3% BSA), followed by addition of 2 ml of Secondary antibody incubation solution per dish, including diluted antibodies according to the table below. Dishes were incubated overnight at 4°C on an orbital shaker.

| **Antibody** | **Cat. No.** | **Dilution** |
| --- | --- | --- |
| Donkey anti Rabbit Alexa 488 | Thermo Fisher, A21206 | 1:750 |
| Donkey anti Mouse Alexa 568 | Thermo Fisher, A10037 | 1:750 |
| Donkey anti Goat Alexa 647 | Thermo Fisher, A21447 | 1:750 |

Secondary antibody solution was removed and wells were washed twice with 1x PBS +0.1% Tween. 2ml of DAPI incubation solution was added per well, consisting of 1x PBS + 0.01% Tween + 2 ug/ml DAPI, and incubated for 10 minutes at room temperature on an orbital shaker. Wells were washed twice with 1x PBS, then 2 ml of 1x PBS + 0.1 sodium azide was added for storage.

For imaging, CPEMs were transferred to glass bottom 8 well chamber slides (Ibidi, 80827). 250 ul of 1x PBS + 90% glycerol (Millipore Sigma, G2025) + 0.1% sodium azide was added to each well of the 8 well chamber slide. The 6 well plate containing CPEMs was slowly swirled so CPEMs cluster in the center and 50 ul was aspirated and added into the 8 well chamber slide wells. The lid was put on the 8 well chamber slide and sealed with clear nail polish (Ted Pella,114-7). Slides were stored overnight at 4°C to let the CPEMs settle before imaging. CPEMs were imaged on a Zeiss 880 confocal, using a 20x long working distance objective. For optical sectioning in the z-axis, images were taken every 3.89 um.

### CPEM image segmentation and processing

Image stacks were loaded into imageJ, and individual CPEMs were cropped from each image stack and saved as separate image sequences per channel. Image sequences per channel imported into Napari for each CPEMs and segmented using StarDist (stardist-napari 2022.12.6). The following settings were used: Model Type: 2D. Pre-trained model: Versatile (fluorescent nuclei). Normalize image yes, percentile low = 1.00, percentile high = 99.80. Probability/Score threshold: 0.6. Overlap threshold: 0.5. Output type: Labels only. Normalization axes: ZYX. Time-lapse labels: Match to previous frame (via overlap). Remaining parts of other CPEMs in the segmentation label layers were erased, so only data for a single complete CPEM was analyzed. For each channel, the segmentation data was analyzed with RegionProps (napari-skimage-regionprops 0.10.1) to extract positions, size and signal intensity of each individually segmented cell and exported to a data table. Data was filtered with the following parameters: Nuclei with mean intensities >400 (Pou5f1), >200 (Cdx2, Foxa1) were included in downstream analysis. Nuclei with areas >400 <16000 were included in downstream analysis. The number and ratio of cells for each region was plotted using ggplot2. Density plots were made using the ggplot2 geom_bin2d option, using 9 bins. For each region, using centroid data, the x/y coordinates were scaled between 0-1, where 0 was the minimum pixel position where the first cell was along the x/y axis, and 1 the maximum position where the last cell was along the x/y axis. In the z-axis, using centroid data, the z-position was normalized to the first image in which cells were detected for each sample. This accounted for small variations in size and crops of CPEM data, and the position in the imaging buffer during acquisition, to better capture the average pattern. Cavity analysis was done in imageJ [83], by plotting the line intensity profile across a center-stack image of DAPI channel per sample. Plotting and clustering was done using PlotTwist [84].

### scRNA-seq preparation and analysis

CPEMs were grown as described before for 72 hours. 1.5 ml of medium was removed and 1.5 ml of 1x PBS was added per well. 1.5 ml of this solution was removed and 0.5 ml of 1x PBS was added per well and CPEMs were transferred to microcentrifuge tubes using cut-off pipette tips. Tubes were centrifuged for 5 minutes at 300 rcf. Supernatant was removed and CPEMs were gently resuspended in 1ml of DMEM/F12 (Thermo Fisher, 11039021), followed by centrifugation for 5 minutes at 300 rcf again. Supernatant was removed and CPEMs were resuspended in 1 ml of Trypsin-EDTA. CPEMs were pipetted up and down 3 times before incubating for 5 minutes at 37°C. CPEMs were pipetted up and down 6 times and 200 ul of ESC qualified FBS was added to each tube, followed by centrifugation for 5 minutes at 300 rcf. Supernatant was removed and cells were resuspended in 0.5 ml standard differentiation medium and transferred to a cryovial. A sample was taken to measure cell viability, samples with viability >80% were used for downstream application. 0.5 ml of 2x Freezing medium, consisting of 85% ESC qualified FBS / 15% DMSO, was added to each cryovial and gently mixed. Cryovials were placed into a cell freezing container and incubated at −80°C overnight, followed by storage in liquid nitrogen (vapor phase) cell storage before single-cell RNAseq preparation. Approximately 1 million cells were frozen per cryovial. An additional vial was frozen from the same batch of CPEMs for each sample. This additional vial was thawed and assessed for cell viability again, before preparing for single-cell RNA-sequencing.

Cryovials were delivered for processing to Novogene Co, San Jose, USA, where samples were prepared following 10X Genomics v3 single-cell 3’ RNA-seq library preparation. 4 samples were prepared, 5,000 cells were captured for each, and parallel sequenced (NovaSeq 6000) with 200M paired read output per sample, resulting in approximately 40,000 paired reads per cell per sample.

Fastq files were aligned and gene counts and barcodes were processed using kallisto-bustools [85] kb_python-0.28.2 with the code below:

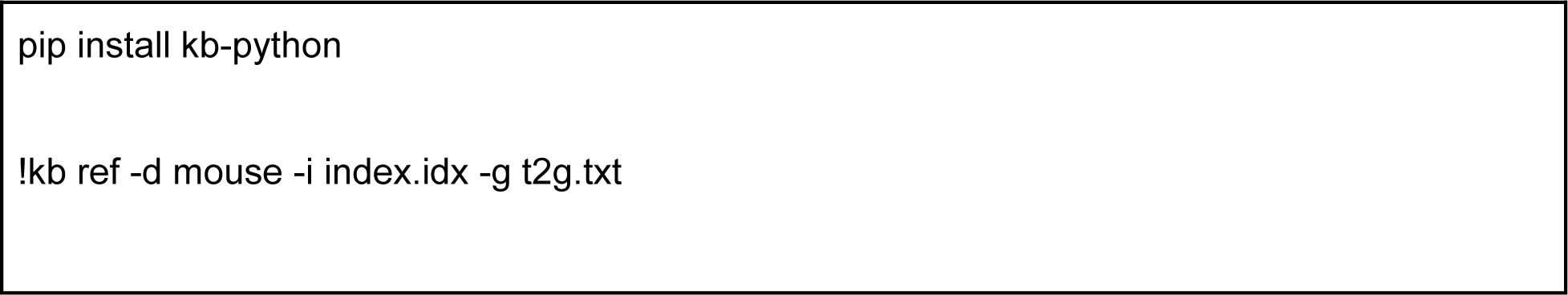
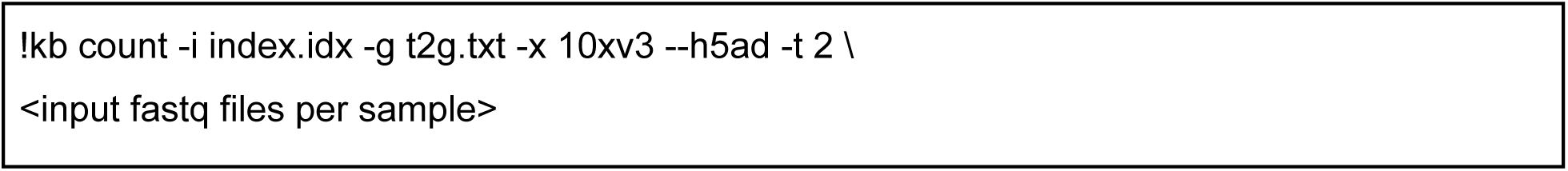

The resulting output matrices were filtered and converted to seurat files (Seurat 5.0.1) [86] with the code below:

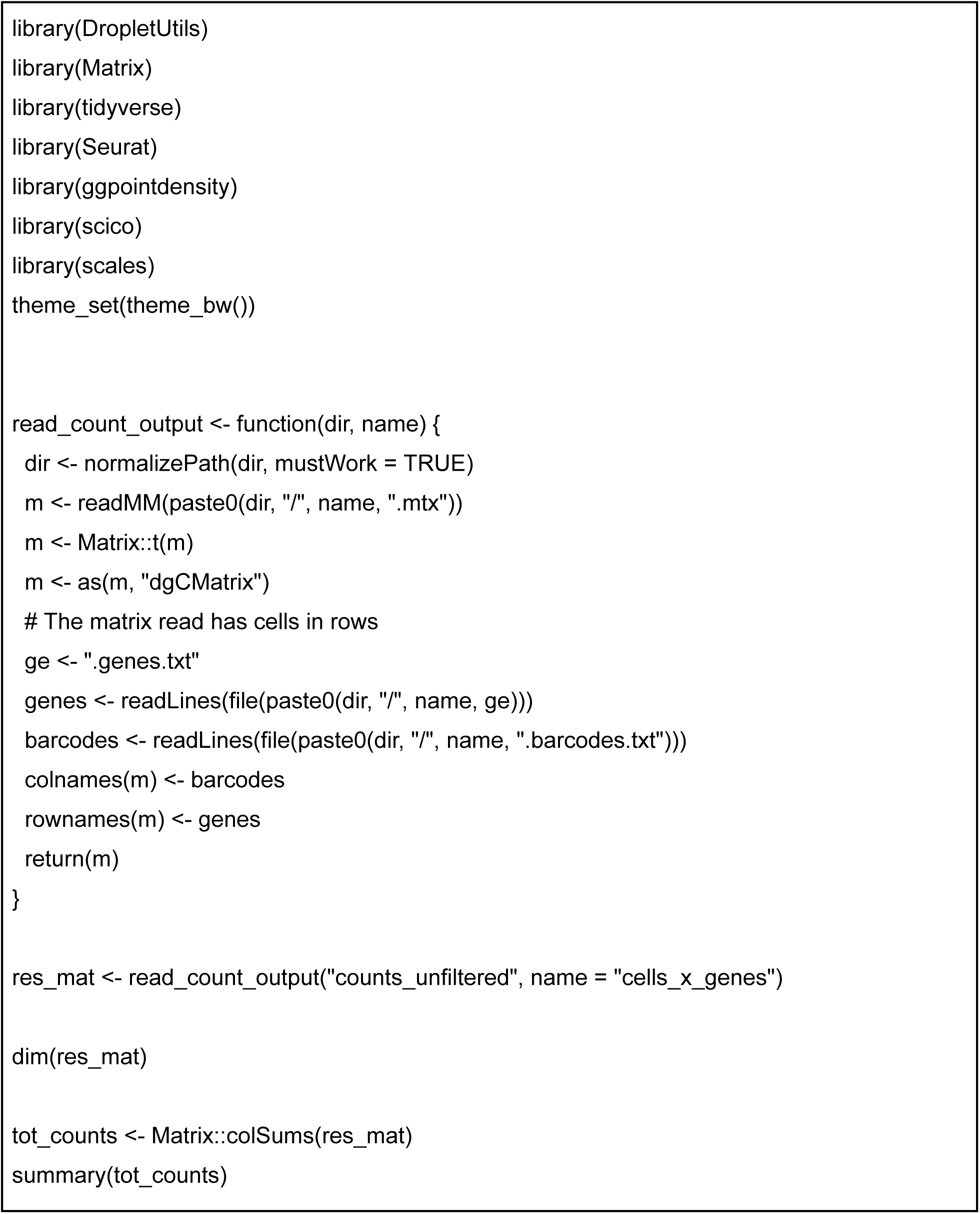
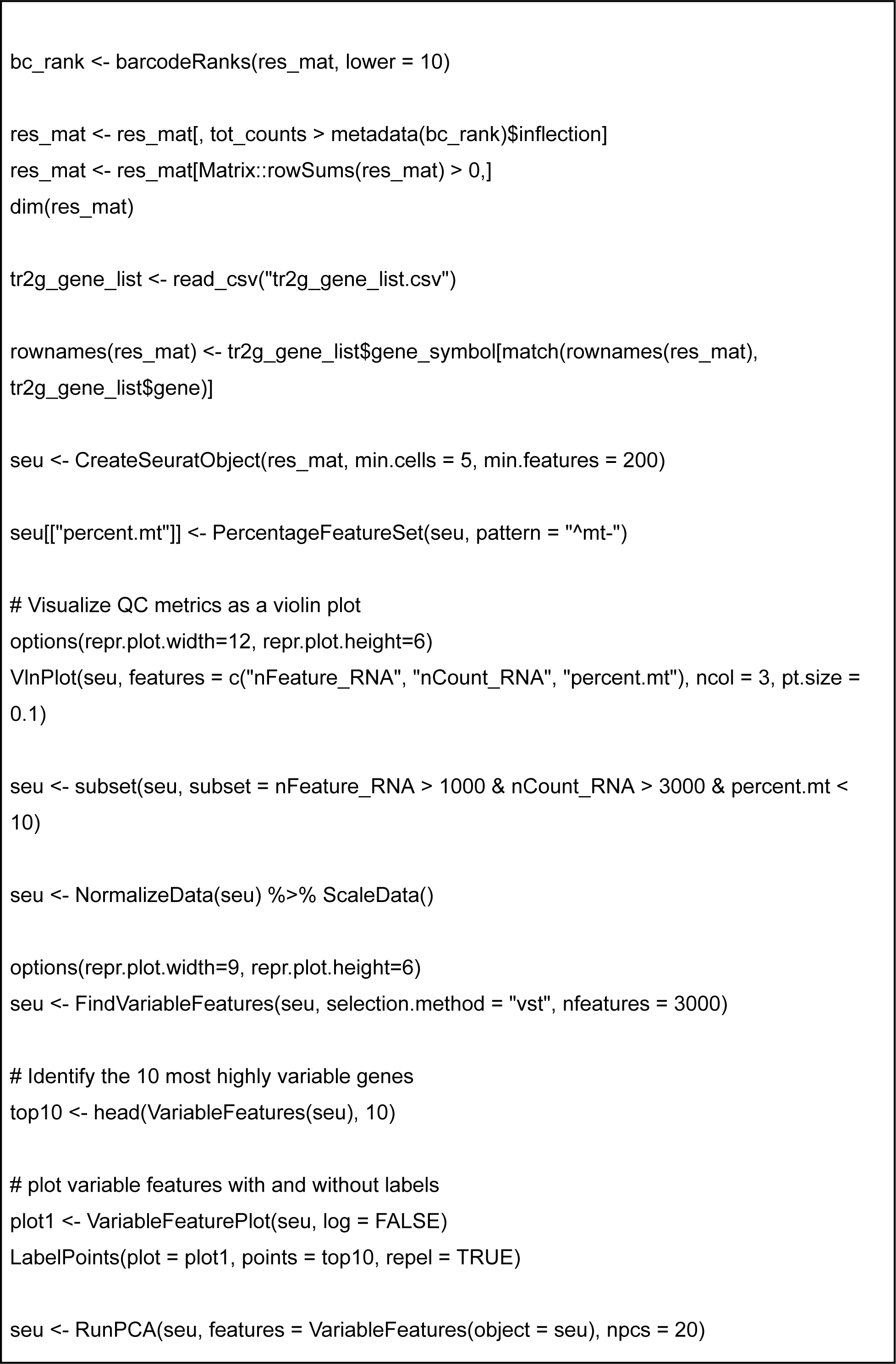
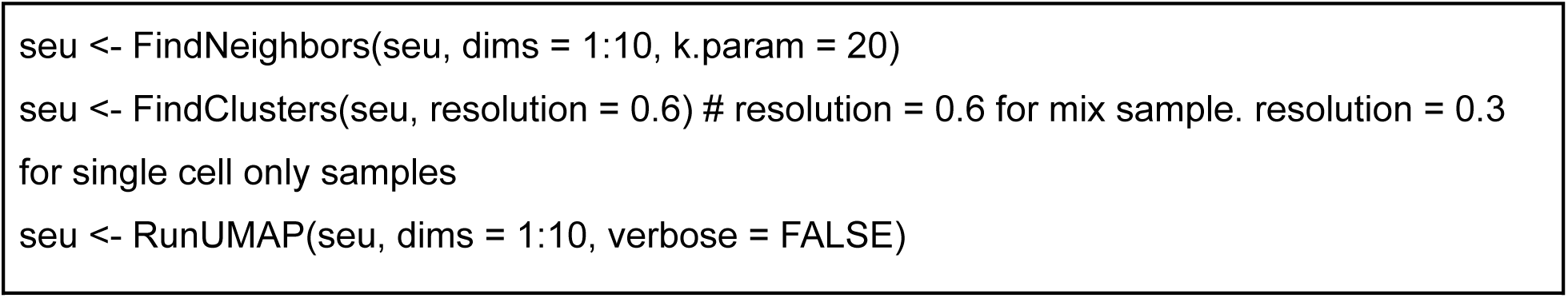

The seurat files were used to prepare umap plots, featureplots and dotplots shown. The seurat for the mixed cell CPEM sample was also used for ligand-receptor interactions using CellChat (2.1.1) with the code below:

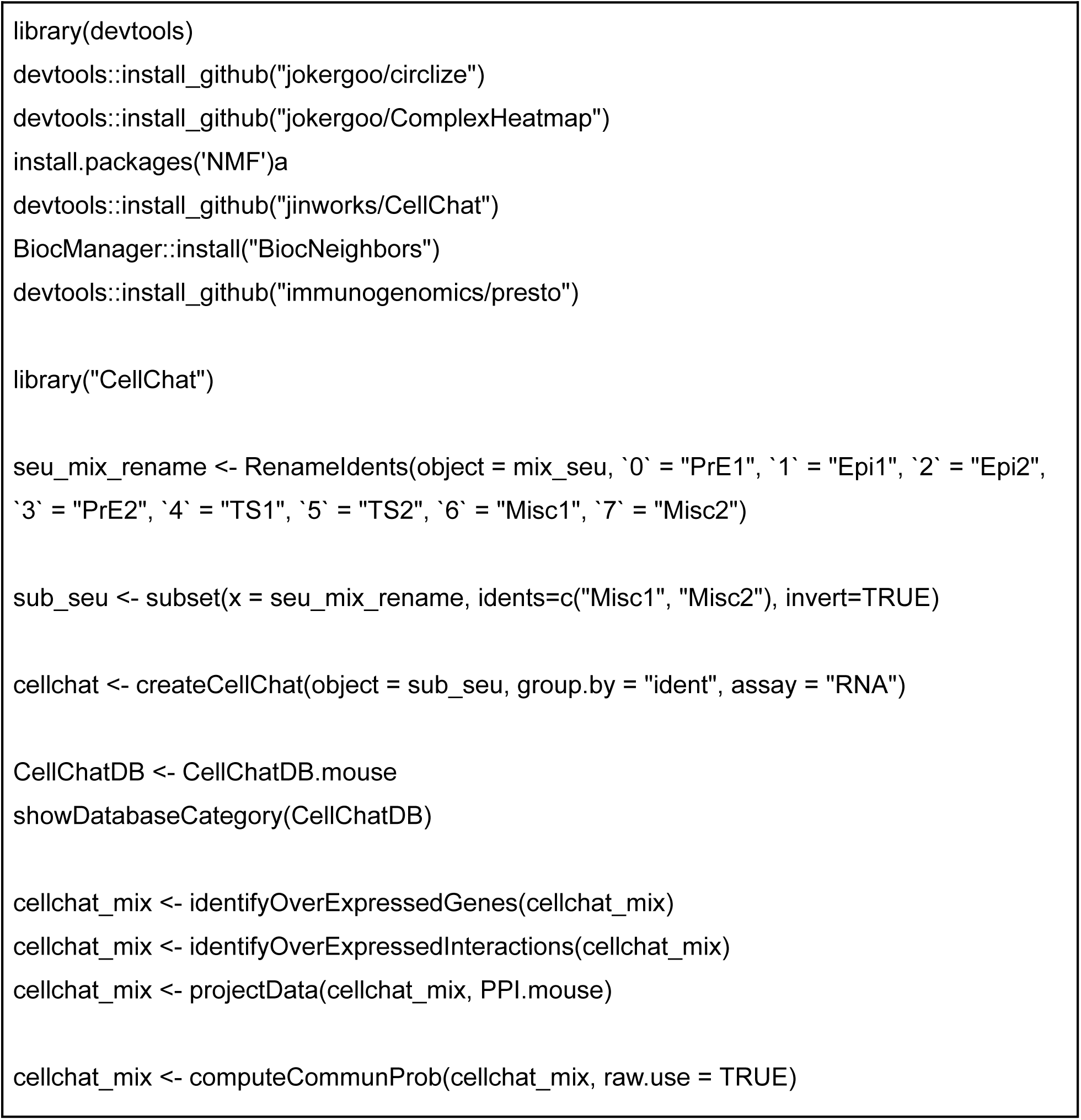
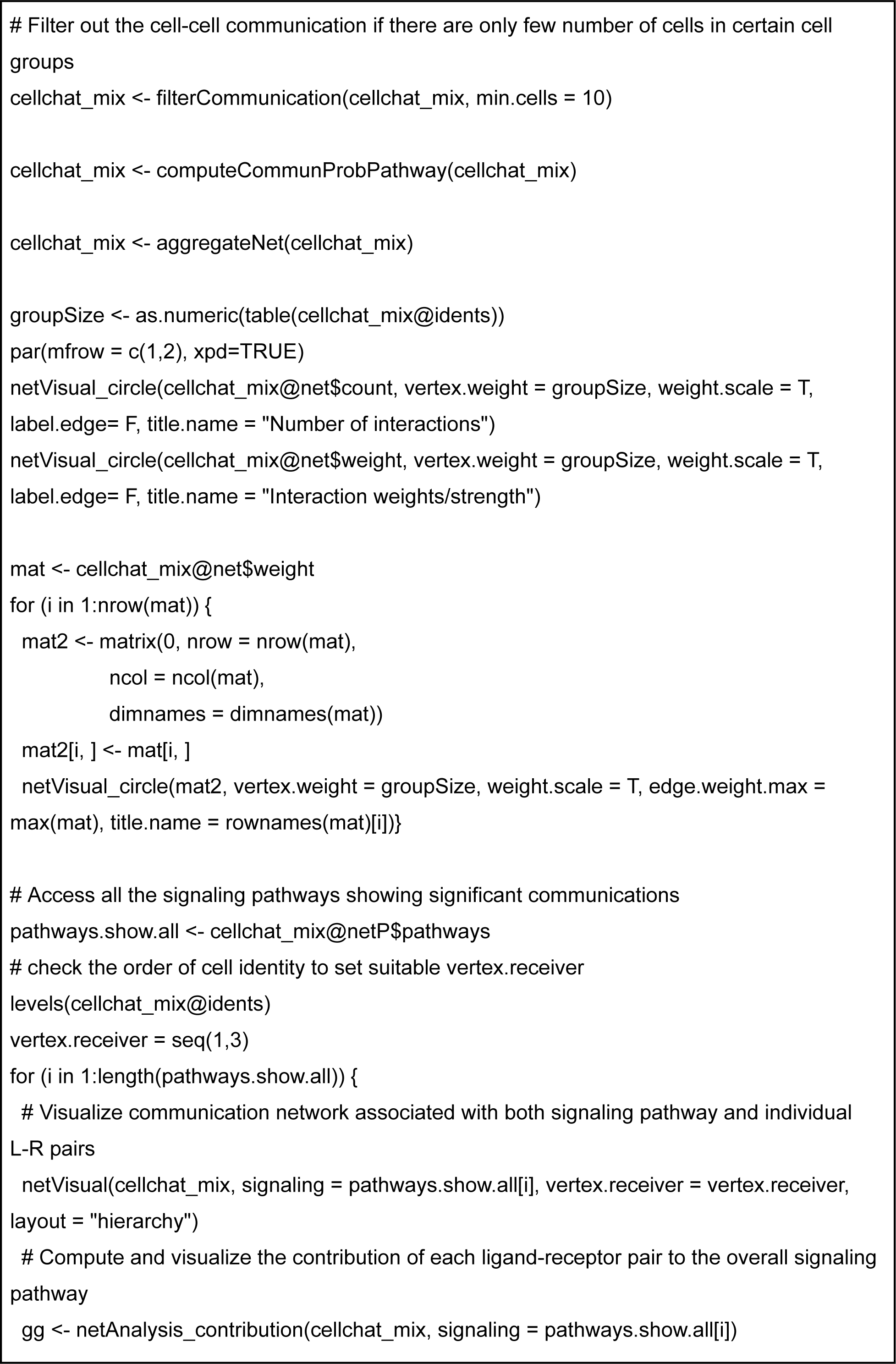
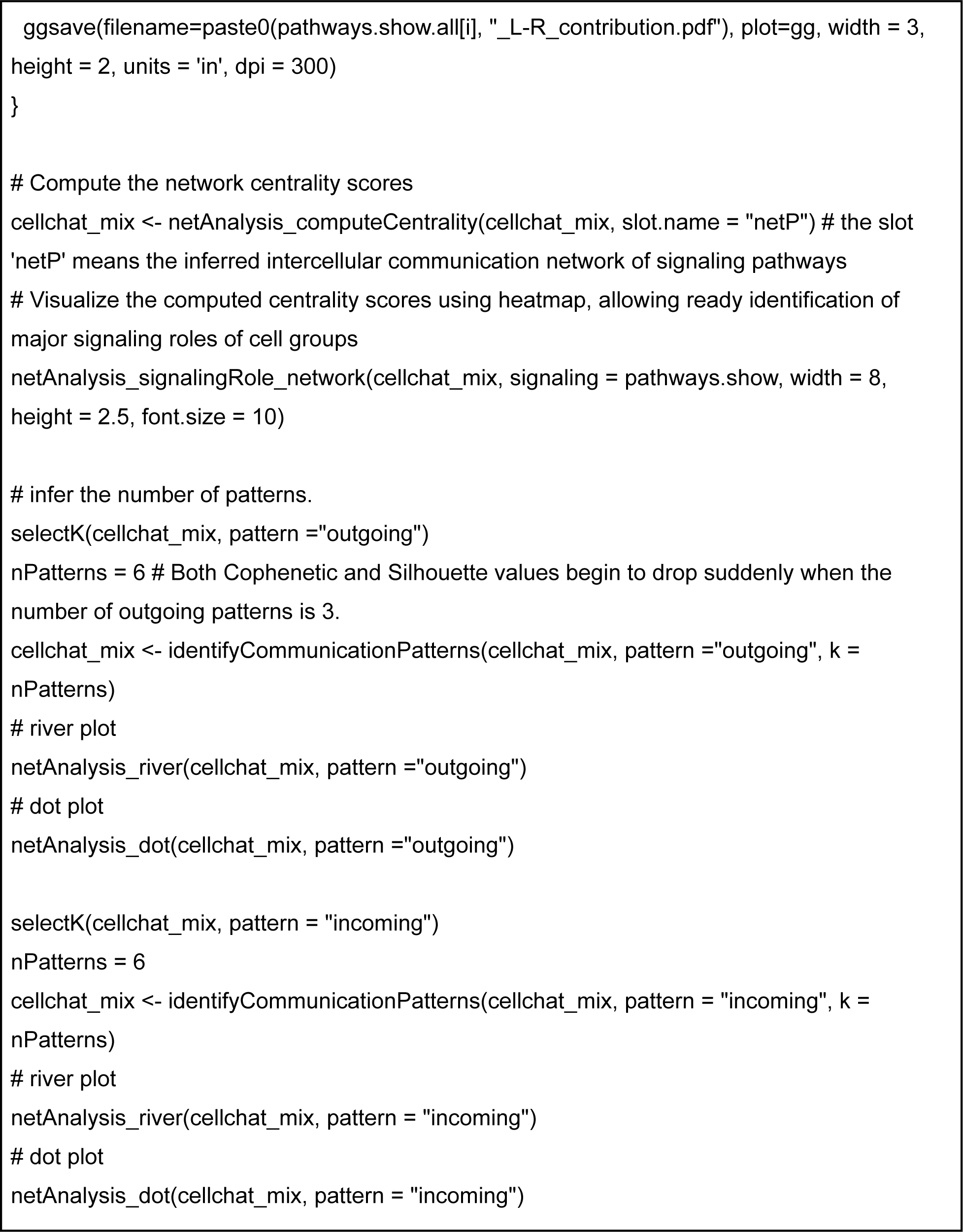

## Data Availability

Raw data, count matrices, seurat files and CellChat file of single-cell RNA-seq analysis are available at GEO under accession number GSE256138.

## Author Contribution

S.A, G.A.L and S.K conceptualized the project, performed experiments and analyzed the majority of the results. S.A and G.A.L wrote the manuscript and prepared the figures. C.H performed immunostaining and analyzed confocal images of the 3D embryo models. A.Z and S.L contributed to the analysis of live imaging 2D micropattern experiments. L.S.Q. provided feedback on the manuscript and CRISPR resources. G.K and R.A fabricated the micropattern dishes used for live imaging experiments. B.R.T set up initial experiments using 2D micropatten culture and reviewed the manuscript.

## Acknowledgements

We would like to thank Dr. Camilla Forsberg, Dr. Russ Corbett-Detig and Dr. Euiseok Kim for providing feedback on our manuscript. We also would like to thank the members of the Shariati lab, in particular Dr. for their feedback on the manuscript.

## References

1. Liu X, Tan JP, Schröder J, Aberkane A, Ouyang JF, Mohenska M, et al. Modelling human blastocysts by reprogramming fibroblasts into iBlastoids. Nature. 2021;591: 627–632.

2. Kagawa H, Javali A, Khoei HH, Sommer TM, Sestini G, Novatchkova M, et al. Human blastoids model blastocyst development and implantation. Nature. 2022;601: 600–605.

3. Weatherbee BAT, Gantner CW, Iwamoto-Stohl LK, Daza RM, Hamazaki N, Shendure J, et al. Pluripotent stem cell-derived model of the post-implantation human embryo. Nature. 2023;622: 584–593.

4. Lau KYC, Rubinstein H, Gantner CW, Hadas R, Amadei G, Stelzer Y, et al. Mouse embryo model derived exclusively from embryonic stem cells undergoes neurulation and heart development. Cell Stem Cell. 2022;29: 1445–1458.e8.

5. Sozen B, Cox AL, De Jonghe J, Bao M, Hollfelder F, Glover DM, et al. Self-Organization of Mouse Stem Cells into an Extended Potential Blastoid. Dev Cell. 2019;51: 698–712.e8.

6. Shahbazi MN, Siggia ED, Zernicka-Goetz M. Self-organization of stem cells into embryos: A window on early mammalian development. Science. 2019;364: 948–951.

7. Rivron NC, Frias-Aldeguer J, Vrij EJ, Boisset J-C, Korving J, Vivié J, et al. Blastocyst-like structures generated solely from stem cells. Nature. 2018;557: 106–111.

8. Warmflash A, Sorre B, Etoc F, Siggia ED, Brivanlou AH. A method to recapitulate early embryonic spatial patterning in human embryonic stem cells. Nat Methods. 2014;11: 847–854.

9. Oldak B, Wildschutz E, Bondarenko V, Comar M-Y, Zhao C, Aguilera-Castrejon A, et al. Complete human day 14 post-implantation embryo models from naive ES cells. Nature. 2023;622: 562–573.

10. Tarazi S, Aguilera-Castrejon A, Joubran C, Ghanem N, Ashouokhi S, Roncato F, et al. Post-gastrulation synthetic embryos generated ex utero from mouse naive ESCs. Cell. 2022;185: 3290–3306.e25.

11. Yanagida A, Spindlow D, Nichols J, Dattani A, Smith A, Guo G. Naive stem cell blastocyst model captures human embryo lineage segregation. Cell Stem Cell. 2021;28: 1016–1022.e4.

12. Yu L, Wei Y, Duan J, Schmitz DA, Sakurai M, Wang L, et al. Blastocyst-like structures generated from human pluripotent stem cells. Nature. 2021;591: 620–626.

13. Sozen B, Jorgensen V, Weatherbee BAT, Chen S, Zhu M, Zernicka-Goetz M. Reconstructing aspects of human embryogenesis with pluripotent stem cells. Nat Commun. 2021;12: 5550.

14. Simunovic M, Metzger JJ, Etoc F, Yoney A, Ruzo A, Martyn I, et al. A 3D model of a human epiblast reveals BMP4-driven symmetry breaking. Nat Cell Biol. 2019;21: 900–910.

15. Amadei G, Handford CE, Qiu C, De Jonghe J, Greenfeld H, Tran M, et al. Embryo model completes gastrulation to neurulation and organogenesis. Nature. 2022;610: 143–153.

16. Zheng Y, Xue X, Shao Y, Wang S, Esfahani SN, Li Z, et al. Controlled modelling of human epiblast and amnion development using stem cells. Nature. 2019;573: 421–425.

17. Moris N, Anlas K, van den Brink SC, Alemany A, Schröder J, Ghimire S, et al. An in vitro model of early anteroposterior organization during human development. Nature. 2020;582: 410–415.

18. Sozen B, Amadei G, Cox A, Wang R, Na E, Czukiewska S, et al. Self-assembly of embryonic and two extra-embryonic stem cell types into gastrulating embryo-like structures. Nat Cell Biol. 2018;20: 979–989.

19. Simunovic M, Siggia ED, Brivanlou AH. In vitro attachment and symmetry breaking of a human embryo model assembled from primed embryonic stem cells. Cell Stem Cell. 2022;29: 962–972.e4.

20. Wei Y, Zhang E, Yu L, Ci B, Sakurai M, Guo L, et al. Dissecting embryonic and extraembryonic lineage crosstalk with stem cell co-culture. Cell. 2023;186: 5859–5875.e24.

21. Hislop J, Song Q, Keshavarz F K, Alavi A, Schoenberger R, LeGraw R, et al. Modelling post-implantation human development to yolk sac blood emergence. Nature. 2024;626: 367–376.

22. Yu L, Logsdon D, Pinzon-Arteaga CA, Duan J, Ezashi T, Wei Y, et al. Large-scale production of human blastoids amenable to modeling blastocyst development and maternal-fetal cross talk. Cell Stem Cell. 2023;30: 1246–1261.e9.

23. Beccari L, Moris N, Girgin M, Turner DA, Baillie-Johnson P, Cossy A-C, et al. Multi-axial self-organization properties of mouse embryonic stem cells into gastruloids. Nature. 2018;562: 272–276.

24. Seong J, Frias-Aldeguer J, Holzmann V, Kagawa H, Sestini G, Heidari Khoei H, et al. Epiblast inducers capture mouse trophectoderm stem cells in vitro and pattern blastoids for implantation in utero. Cell Stem Cell. 2022;29: 1102–1118.e8.

25. Pedroza M, Gassaloglu SI, Dias N, Zhong L, Hou T-CJ, Kretzmer H, et al. Self-patterning of human stem cells into post-implantation lineages. Nature. 2023;622: 574–583.

26. Shahbazi MN, Zernicka-Goetz M. Deconstructing and reconstructing the mouse and human early embryo. Nat Cell Biol. 2018;20: 878–887.

27. Liu P, Chen M, Liu Y, Qi LS, Ding S. CRISPR-Based Chromatin Remodeling of the Endogenous Oct4 or Sox2 Locus Enables Reprogramming to Pluripotency. Cell Stem Cell. 2018;22: 252–261.e4.

28. Liu Y, Yu C, Daley TP, Wang F, Cao WS, Bhate S, et al. CRISPR Activation Screens Systematically Identify Factors that Drive Neuronal Fate and Reprogramming. Cell Stem Cell. 2018;23: 758–771.e8.

29. Pulecio J, Verma N, Mejía-Ramírez E, Huangfu D, Raya A. CRISPR/Cas9-Based Engineering of the Epigenome. Cell Stem Cell. 2017;21: 431–447.

30. Sokka J, Yoshihara M, Kvist J, Laiho L, Warren A, Stadelmann C, et al. CRISPR activation enables high-fidelity reprogramming into human pluripotent stem cells. Stem Cell Reports. 2022;17: 413–426.

31. Ralston A, Rossant J. Cdx2 acts downstream of cell polarization to cell-autonomously promote trophectoderm fate in the early mouse embryo. Dev Biol. 2008;313: 614–629.

32. Ralston A, Cox BJ, Nishioka N, Sasaki H, Chea E, Rugg-Gunn P, et al. Gata3 regulates trophoblast development downstream of Tead4 and in parallel to Cdx2. Development. 2010;137: 395–403.

33. Schrode N, Saiz N, Di Talia S, Hadjantonakis A-K. GATA6 levels modulate primitive endoderm cell fate choice and timing in the mouse blastocyst. Dev Cell. 2014;29: 454–467.

34. Wamaitha SE, del Valle I, Cho LTY, Wei Y, Fogarty NME, Blakeley P, et al. Gata6 potently initiates reprograming of pluripotent and differentiated cells to extraembryonic endoderm stem cells. Genes Dev. 2015;29: 1239–1255.

35. Chowdhary S, Hadjantonakis A-K. Journey of the mouse primitive endoderm: from specification to maturation. Philos Trans R Soc Lond B Biol Sci. 2022;377: 20210252.

36. Qiao Y, Ren C, Huang S, Yuan J, Liu X, Fan J, et al. High-resolution annotation of the mouse preimplantation embryo transcriptome using long-read sequencing. Nat Commun. 2020;11: 2653.

37. Chromatin Accessibility Landscape in Human Early Embryos and Its Association with Evolution. Cell. 2018;173: 248–259.e15.

38. Morgani SM, Metzger JJ, Nichols J, Siggia ED, Hadjantonakis A-K. Micropattern differentiation of mouse pluripotent stem cells recapitulates embryo regionalized cell fate patterning. Elife. 2018;7. doi:10.7554/eLife.32839

39. Knight GT, Lundin BF, Iyer N, Ashton LM, Sethares WA, Willett RM, et al. Engineering induction of singular neural rosette emergence within hPSC-derived tissues. Elife. 2018;7. doi:10.7554/eLife.37549

40. Ahlers J, Althviz Moré D, Amsalem O, Anderson A, Bokota G, Boone P, et al. napari: a multi-dimensional image viewer for Python. Zenodo; 2023. doi:10.5281/ZENODO.3555620

41. Ulicna K, Vallardi G, Charras G, Lowe AR. Automated Deep Lineage Tree Analysis Using a Bayesian Single Cell Tracking Approach. Front Comput Sci. 2021;3: 734559.

42. Schmidt U, Weigert M, Broaddus C, Myers G. Cell detection with star-convex polygons. Medical Image Computing and Computer Assisted Intervention – MICCAI 2018. Cham: Springer International Publishing; 2018. pp. 265–273.

43. Weigert M, Schmidt U, Haase R, Sugawara K, Myers G. Star-convex polyhedra for 3D object detection and segmentation in microscopy. 2020 IEEE Winter Conference on Applications of Computer Vision (WACV). IEEE; 2020. doi:10.1109/wacv45572.2020.9093435

44. Zargari A, Lodewijk GA, Mashhadi N, Cook N, Neudorf CW, Araghbidikashani K, et al. DeepSea is an efficient deep-learning model for single-cell segmentation and tracking in time-lapse microscopy. Cell Rep Methods. 2023;3: 100500.

45. Moghe P, Belousov R, Ichikawa T, Iwatani C, Tsukiyama T, Graner F, et al. Apical-driven cell sorting optimised for tissue geometry ensures robust patterning. bioRxiv. 2023. p. 2023.05.16.540918. doi:10.1101/2023.05.16.540918

46. Amadei G, Lau KYC, De Jonghe J, Gantner CW, Sozen B, Chan C, et al. Inducible Stem-Cell-Derived Embryos Capture Mouse Morphogenetic Events In Vitro. Dev Cell. 2021;56: 366–382.e9.

47. Dupont C, Schäffers OJM, Tan BF, Merzouk S, Bindels EM, Zwijsen A, et al. Efficient generation of ETX embryoids that recapitulate the entire window of murine egg cylinder development. Sci Adv. 2023;9: eadd2913.

48. Klumpe HE, Langley MA, Linton JM, Su CJ, Antebi YE, Elowitz MB. The context-dependent, combinatorial logic of BMP signaling. Cell systems. 2022. pp. 388–407.e10.

49. Pham PD, Lu H, Han H, Zhou JJ, Madan A, Wang W, et al. Transcriptional network governing extraembryonic endoderm cell fate choice. Dev Biol. 2023;502: 20–37.

50. Latos PA, Sienerth AR, Murray A, Senner CE, Muto M, Ikawa M, et al. Elf5-centered transcription factor hub controls trophoblast stem cell self-renewal and differentiation through stoichiometry-sensitive shifts in target gene networks. Genes Dev. 2015;29: 2435–2448.

51. Lin S-CJ, Wani MA, Whitsett JA, Wells JM. Klf5 regulates lineage formation in the pre-implantation mouse embryo. Development. 2010;137: 3953–3963.

52. James JL, Hurley DG, Gamage TKJB, Zhang T, Vather R, Pantham P, et al. Isolation and characterisation of a novel trophoblast side-population from first trimester placentae. Reproduction. 2015;150: 449–462.

53. Huang D, Guo G, Yuan P, Ralston A, Sun L, Huss M, et al. The role of Cdx2 as a lineage specific transcriptional repressor for pluripotent network during the first developmental cell lineage segregation. Sci Rep. 2017;7: 17156.

54. Qiu C, Cao J, Martin BK, Li T, Welsh IC, Srivatsan S, et al. Systematic reconstruction of cellular trajectories across mouse embryogenesis. Nat Genet. 2022;54: 328–341.

55. Okubo T, Rivron N, Kabata M, Masaki H, Kishimoto K, Semi K, et al. Hypoblast from human pluripotent stem cells regulates epiblast development. Nature. 2024;626: 357–366.

56. Chandrasekaran AP, Li M. Extra (embryonic) dialogues: Keys to improved stem cell-based embryo models. Cell Stem Cell. 2024;31: 155–157.

57. Jin S, Plikus MV, Nie Q. CellChat for systematic analysis of cell-cell communication from single-cell and spatially resolved transcriptomics. bioRxiv. 2023. p. 2023.11.05.565674. doi:10.1101/2023.11.05.565674

58. Jin S, Guerrero-Juarez CF, Zhang L, Chang I, Ramos R, Kuan C-H, et al. Inference and analysis of cell-cell communication using CellChat. Nat Commun. 2021;12: 1–20.

59. Goissis MD, Bradshaw B, Posfai E, Rossant J. Influence of FGF4 and BMP4 on FGFR2 dynamics during the segregation of epiblast and primitive endoderm cells in the pre-implantation mouse embryo. PLoS One. 2023;18: e0279515.

60. Christodoulou N, Weberling A, Strathdee D, Anderson KI, Timpson P, Zernicka-Goetz M. Morphogenesis of extra-embryonic tissues directs the remodelling of the mouse embryo at implantation. Nat Commun. 2019;10: 3557.

61. Molotkov A, Soriano P. Distinct mechanisms for PDGF and FGF signaling in primitive endoderm development. Dev Biol. 2018;442: 155–161.

62. Graham SJL, Wicher KB, Jedrusik A, Guo G, Herath W, Robson P, et al. BMP signalling regulates the pre-implantation development of extra-embryonic cell lineages in the mouse embryo. Nat Commun. 2014;5: 1–11.

63. Simon CS, Rahman S, Raina D, Schröter C, Hadjantonakis A-K. Live Visualization of ERK Activity in the Mouse Blastocyst Reveals Lineage-Specific Signaling Dynamics. Dev Cell. 2020;55: 341–353.e5.

64. Artus J, Panthier J-J, Hadjantonakis A-K. A role for PDGF signaling in expansion of the extra-embryonic endoderm lineage of the mouse blastocyst. Development. 2010;137: 3361–3372.

65. Cang Z, Wang Y, Wang Q, Cho KWY, Holmes W, Nie Q. A multiscale model via single-cell transcriptomics reveals robust patterning mechanisms during early mammalian embryo development. PLoS Comput Biol. 2021;17: e1008571.

66. Maye P, Becker S, Kasameyer E, Byrd N, Grabel L. Indian hedgehog signaling in extraembryonic endoderm and ectoderm differentiation in ES embryoid bodies. Mech Dev. 2000;94: 117–132.

67. Yuan J, Cha J, Deng W, Bartos A, Sun X, Ho H-YH, et al. Planar cell polarity signaling in the uterus directs appropriate positioning of the crypt for embryo implantation. Proc Natl Acad Sci U S A. 2016;113: E8079–E8088.

68. Fan R, Kim YS, Wu J, Chen R, Zeuschner D, Mildner K, et al. Wnt/Beta-catenin/Esrrb signalling controls the tissue-scale reorganization and maintenance of the pluripotent lineage during murine embryonic diapause. Nat Commun. 2020;11: 5499.

69. Semaphorin-Plexin Signaling Controls Mitotic Spindle Orientation during Epithelial Morphogenesis and Repair. Dev Cell. 2015;33: 299–313.

70. Rugg-Gunn PJ, Cox BJ, Lanner F, Sharma P, Ignatchenko V, McDonald ACH, et al. Cell-surface proteomics identifies lineage-specific markers of embryo-derived stem cells. Dev Cell. 2012;22: 887–901.

71. Bao M, Cornwall-Scoones J, Sanchez-Vasquez E, Cox AL, Chen D-Y, De Jonghe J, et al. Stem cell-derived synthetic embryos self-assemble by exploiting cadherin codes and cortical tension. Nat Cell Biol. 2022;24: 1341–1349.

72. Saiz N, Mora-Bitria L, Rahman S, George H, Herder JP, Garcia-Ojalvo J, et al. Growth-factor-mediated coupling between lineage size and cell fate choice underlies robustness of mammalian development. Elife. 2020;9. doi:10.7554/eLife.56079

73. Rugg-Gunn PJ, Moris N, Tam PPL. Technical challenges of studying early human development. Development. 2023;150: dev201797.

74. Magnusson JP, Rios AR, Wu L, Qi LS. Enhanced Cas12a multi-gene regulation using a CRISPR array separator. Elife. 2021;10. doi:10.7554/eLife.66406

75. Kempton HR, Goudy LE, Love KS, Qi LS. Multiple Input Sensing and Signal Integration Using a Split Cas12a System. Mol Cell. 2020;78: 184–191.e3.

76. Jiang X, Wang Y, Xiao Z, Yan L, Guo S, Wang Y, et al. A differentiation roadmap of murine placentation at single-cell resolution. Cell Discov. 2023;9: 30.

77. Shariati SA, Dominguez A, Xie S, Wernig M, Qi LS, Skotheim JM. Reversible Disruption of Specific Transcription Factor-DNA Interactions Using CRISPR/Cas9. Mol Cell. 2019;74: 622–633.e4.

78. Truong VA, Hsu M-N, Kieu Nguyen NT, Lin M-W, Shen C-C, Lin C-Y, et al. CRISPRai for simultaneous gene activation and inhibition to promote stem cell chondrogenesis and calvarial bone regeneration. Nucleic Acids Res. 2019;47: e74.

79. Gao Y, Xiong X, Wong S, Charles EJ, Lim WA, Qi LS. Complex transcriptional modulation with orthogonal and inducible dCas9 regulators. Nat Methods. 2016;13: 1043–1049.

80. Concordet J-P, Haeussler M. CRISPOR: intuitive guide selection for CRISPR/Cas9 genome editing experiments and screens. Nucleic Acids Res. 2018;46: W242–W245.

81. Wickham H. ggplot2: Elegant Graphics for Data Analysis. Springer; 2016.

82. Hosmer DW Jr, Lemeshow S, May S. Applied Survival Analysis: Regression Modeling of Time-to-Event Data. John Wiley & Sons; 2011.

83. Schindelin J, Arganda-Carreras I, Frise E, Kaynig V, Longair M, Pietzsch T, et al. Fiji: an open-source platform for biological-image analysis. Nat Methods. 2012;9: 676–682.

84. Goedhart J. PlotTwist: A web app for plotting and annotating continuous data. PLoS Biol. 2020;18: e3000581.

85. Melsted P, Booeshaghi AS, Liu L, Gao F, Lu L, Min KHJ, et al. Modular, efficient and constant-memory single-cell RNA-seq preprocessing. Nat Biotechnol. 2021;39: 813–818.

86. Hao Y, Stuart T, Kowalski MH, Choudhary S, Hoffman P, Hartman A, et al. Dictionary learning for integrative, multimodal and scalable single-cell analysis. Nat Biotechnol. 2024;42: 293–304.

